# Elongasome Dysfunction Triggers Dependence on MepM-Mediated Peptidoglycan Recycling

**DOI:** 10.64898/2026.04.14.718437

**Authors:** Lama Shamseddine, Manon Janet-Maitre, Asmaa Yousra Chikhi, Harinderbir Kaur, Jean-Luc Pellequer, Ina Attrée, Viviana Job

## Abstract

The bacterial peptidoglycan (PG) layer is responsible for maintaining cell shape and protecting the cell against both external stress and internal turgor pressure. In rod-shaped bacteria, PG synthesis is orchestrated by two major multi-protein complexes: the elongasome and the divisome, which drive lateral and septal cell wall synthesis, respectively. The membrane protein MreD plays a central role in regulating the elongasome activity. Here, we leveraged a viable *Pseudomonas aeruginosa* transposon mutant, with reduced *mreD* expression, to dissect cellular adaptations to impaired elongasome function. A genome-wide synthetic lethality screen revealed that cell division and PG recycling pathways become essential in the context of an impaired elongasome machinery. Moreover, Tn-seq analysis identified the endopeptidase MepM as essential in the *mreD* downregulated background, whereas its overexpression promoted cell elongation in the parental strain, supporting a conserved role in lateral wall biogenesis. We demonstrate that MepM sustains viability during elongasome impairment by maintaining an active MurU-dependent PG recycling pathway. Together, our findings uncover a functional coupling between PG hydrolysis, recycling, and synthesis. We also identify MepM and PG recycling as critical determinants of bacterial survival when elongasome activity is compromised, positioning these pathways as attractive targets for the development of next-generation or combinatory antibacterial therapies.

## Introduction

*Pseudomonas aeruginosa* is a rod-shaped opportunistic pathogen that can cause life-threatening infections and exhibits intrinsic resistance to a wide range of antibiotics, including ꞵ-lactams that target bacterial peptidoglycan (PG) synthesis (1). The PG is a key structural component of the cell envelope, which determines bacterial shape and provides protection against the external environment and internal turgor pressure (2).

The PG is assembled from the precursor lipid II, which is produced in the cytoplasm and then flipped into the periplasm (3). Lipid II is a disaccharide-pentapeptide composed of two sugar units, N-acetylglucosamine (GlcNAc) and N-acetylmuramic acid (MurNAc), to which a stem peptide is covalently attached (4). In the periplasm, lipid II is incorporated into the growing cell wall through β-1→4 glycosidic bonds by the glycosyltransferases (GTases) (5) and the stem is crosslinked to existing PG stem by the transpeptidase (TPase) activity of penicillin-binding-proteins (PBPs). In Gram-negative bacteria, this occurs between the fourth D-alanine of a donor stem and the third meso-diaminopimelic acid (mDAP) of an acceptor stem (6).

The PG sacculus is continuously remodeled during bacterial growth through the coordinated action of numerous enzymes (such as endopeptidases, carboxypeptidases, and others) (7). During this process, cleavage of peptidoglycan strands releases soluble muropeptide fragments that can be transported back into the cytoplasm and recycled (8). This recycling pathway allows recovery of both peptide and sugar components, which can be reused for the synthesis of new peptidoglycan precursors (9). PG recycling therefore helps maintain the pool of cell wall building blocks and contributes to cell envelope homeostasis. In parallel, peptidoglycan synthesis during growth is carried out by two machineries, the divisome and the elongasome (Rod system), responsible for the synthesis of the septal and lateral wall, respectively (7).

Although classified as two distinct complexes, both machineries comprise proteins with analogous functions: a cytoskeletal element that organizes the complexes, a Shape, Elongation, Division, Sporulation (SEDS) protein and a monofunctional class B PBP (bPBP) carrying a TPase activity. Together, the SEDS protein and bPBP, form the core “PG synthase” (10). In addition, a bifunctional class A PBP (aPBP) associates with the complex and contributes to both GTase and TPase activities (10,11).

In the divisome, the organization relies on the tubulin-like protein FtsZ, which assembles into filament bundles at mid-cell (12). The SEDS protein FtsW partners with PBP3 to form the septal PG synthase, while the bifunctional PBP1b, associates with the divisome and contributes to cell wall polymerization (13). Similarly, the elongasome relies on the actin-like protein MreB, forming anti-parallel filaments that guide the spatial and functional organization of this machinery (Figure 1A) (14). Here, the core elongasome PG synthase consisting of PBP2 and the SEDS protein RodA, act together with PBP1a to mediate lateral wall PG synthesis (15).

**Figure 1.**
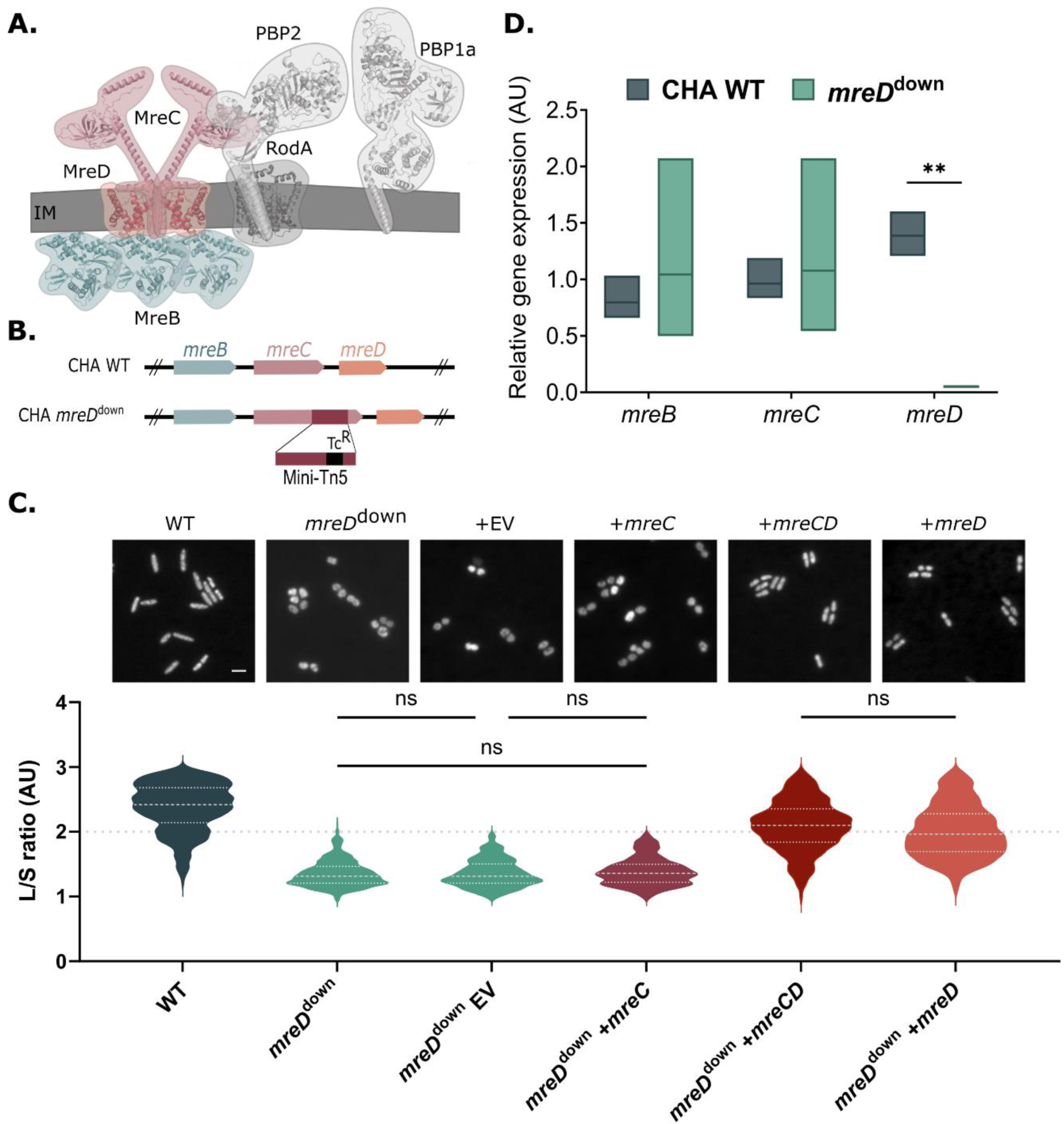
Loss of rod shape in the *mreC* transposon mutant is due to decreased expression of *mreD*. **A.** Scheme of the elongasome machinery in rod-shaped bacteria. MreB forms filaments in close proximity to the inner membrane (IM) and recruits elongasome proteins. Peptidoglycan synthesis requires an aPBP (PBP1a), a peptidoglycan synthase consisting of PBP2 and RodA, and the regulatory proteins MreC and MreD. High-resolution structures of individual elongasome components are shown in transparency, consistent with published complexes: PBP2/MreC (PDB 5lp5), PBP2-TM/RodA (PDB 6pl5), MreC-TM/MreD (PDB 9dvb) and MreB oligomers (PDB 1JCG). PBP1a and MreC-CTD from PAO1 are presented as AlphaFold3 models. **B.** Schematic representation of transposon insertion in *mreD^down^*. **C.** Morphological characterization of wild-type (WT) and *mreD^down^* cells. An empty vector pJN105 (EV) or a pJN105 harboring *mreC*, *mreCD* or *mreD* was introduced into the transposon mutant *mreD^down^*. Bacteria were labeled with Syto24 DNA dye and images were obtained using fluorescence microscopy. Scale: 1 µm. Bacterial morphology was analyzed using the MicrobeJ plugin of ImageJ (28). L/S ratio represents the ratio of the long versus the short axis. n=450 bacteria. A Kruskal-Wallis test was performed and all groups had a statistically significant difference (*p-val* < 0.001) except for the comparisons marked non-significant (ns). The three visible lines represent the median and the 1^st^ and 3^rd^ quartiles. **D.** Relative gene expression of *mreBCD* operon analyzed by RT-qPCR in the wild-type and the transposon mutant *mreD^down^*. For each gene tested, a Welch’s t-test was performed. The line in the boxplot shows the mean.

Two additional proteins, MreC and MreD, are part of the core elongasome complex (Figure 1A) (16). While MreD is confined to the inner membrane, MreC is a transmembrane protein that extends into the periplasm. MreD interacts with MreC within the bilayer, allowing a tilting motion of the periplasmic β-domain of MreC, which extends above the bacterial inner membrane (17). This reorientation allows MreC to interact with the non-catalytic periplasmic pedestal domain of PBP2, triggering its opening and elevating the active site of PBP2 towards the PG layers (17–19). When activated, the PBP2 pedestal domain is thought to be the key allosteric activator of its interacting partner RodA (19,20).

MreB, MreC and MreD are encoded by the *mreBCD* operon (21) (Figure 1B). In *Bacillus subtilis* and *Escherichia coli,* the deletion of *mreB* results in non-viable, enlarged spherical cells, reflecting the essentiality of *mreB* in maintaining bacterial rod-shape and cell integrity (22,23). In addition, depletion of MreC or MreD abolishes the formation of MreB filaments in *E. coli*, highlighting the structural interdependence of these proteins within the Rod system (22). Despite the central role of the Rod system in bacterial morphogenesis, how rod bacteria adapt to defects in elongasome activity remains poorly understood. In most rod-shaped bacteria, deletion of genes within the *mreBCD* operon is lethal, which has limited the investigation of the physiological consequences of Rod system loss and the cellular pathways that compensate for it. In particular, the extent to which perturbations in lateral PG synthesis affect other processes involved in cell envelope homeostasis, such as septal PG synthesis and PG recycling, remains largely unexplored. Addressing these questions is particularly relevant in opportunistic pathogens such as *Pseudomonas aeruginosa*, which displays intrinsic resistance to many antibiotics targeting PG synthesis (24). Its robust cell envelope, therefore, makes it a valuable model to study how bacteria maintain envelope integrity under cell wall stress.

In this work, we used a viable mutant of *P. aeruginosa* carrying a transposon insertion in the *mreC* gene. We show that the loss of rod shape in this mutant is due to a downregulation of *mreD*. In this context, the impaired elongasome activity is compensated by increased divisome activity. To further investigate the mechanisms that allow survival under elongasome dysfunction, we performed a synthetic lethality screen to identify conditionally essential genes in this background. This approach revealed connections between PG synthesis and PG recycling, suggesting that recycling-derived precursors contribute to maintaining cell wall homeostasis when elongation is compromised. Notably, our results highlight the importance of MepM endopeptidase in sustaining the PG precursor pool independently of *de novo* PG synthesis in this mutant. Together, these findings uncover adaptive pathways that support cell envelope integrity when elongasome function is impaired and reveal potential vulnerabilities that could be exploited for antimicrobial strategies.

## Results

### Reduced expression of *mreD* is responsible for the loss of cell shape in *mreD^down^* mutant

We took advantage of a viable mutant harboring a Tn5-Tet^R^ transposon insertion in *mreC* in the clinical strain CHA (Figure 1B) to investigate the effect of elongasome dysfunction in *P. aeruginosa* (25). Insertion of the transposon at the 3’ end of the gene resulted in the production of a hybrid MreC^Tn^ protein of 35.4 kDa with a modified C-terminal sequence compared to 35.2 kDa wild-type MreC (MreC^WT^) (Figure 1B and S1A). MreC^Tn^ protein levels were reduced compared to MreC^WT^, and we observed the presence of a degradation product(s) during bacterial growth (Figure S1B). As previously reported for strains with diminished MreC levels, the transposon insertion in *mreC* led to spherical cells, in contrast to the rod-shaped parental strain (Figure 1C) (26,27).

However, the supplementation of the transposon mutant with a wild-type copy of *mreC* did not restore rod shape, indicating that alteration of the MreC CTD alone was not responsible for the spherical phenotype (Figure 1C). In contrast, rod shape was restored upon *in trans* addition of *mreD*, either alone or in combination with *mreC* (Figure 1C). This is consistent with RT-qPCR analysis showing a 30-fold decrease in *mreD* expression in the transposon mutant, whereas *mreC* expression remained unchanged (Figure 1D). Together, our data show that downregulation of *mreD* in the transposon mutant (hereafter referred to as *mreD^down^*), leading to the impaired elongasome, is responsible for the loss of rod shape.

### Major phenotypic defects due to impaired elongasome

To characterize the consequences of an impaired elongasome on *P. aeruginosa* biology, we used the *mreD^down^* strain, which exhibits no obvious growth defects in rich medium (Figure S2A), and examined swimming, cytotoxicity, and alginate production. *P. aeruginosa* is a monotrichous bacterium whose single polar flagellum is required for swimming in liquid environments (29,30). Swimming motility was strongly impaired in the *mreD^down^* strain compared to the parental strain (Figure S2B), suggesting reduced expression of genes involved in flagellum biosynthesis. Unexpectedly, transcriptomic analysis during exponential growth revealed overexpression of most genes from this pathway in the *mreD^down^* strain (Figure S2C; Table S1), while the expression of *flhF* and *fleN*, which control flagellar number and polar positioning, remained unchanged (Figure S2C) (29,31). These results suggest a defect in flagellar assembly/anchoring rather than flagellar biogenesis. This hypothesis was supported by immunofluorescence labeling of the flagellar structural protein FliC. Although flagellated cells were detected in both the wild-type and *mreD^down^* samples, detached flagella were frequently observed in the latter (Figure S2D). Whereas *mreD*^down^ cells are capable of producing and assembling flagella, our results suggest that anchoring of the flagellar structure to the bacterial surface is compromised. To determine whether this structural defect extends to other virulence-associated systems, we next examined the Type III Secretion System (T3SS) in the mutant. The T3SS is a major virulence factor of *P. aeruginosa,* used as a molecular syringe to inject effector proteins into host cells and induce cytotoxicity (32). Both the T3SS structural components PscF (needle filament) and PcrV (needle-pore bridge) were present at the surface of *mreD^down^* (Figure S2E), and the mutant retained cytotoxicity toward J774A.1 murine macrophages comparable to the parental strain, consistent with a functional T3SS (Figure S2F).

Moreover, given that the parental strain is highly mucoid due to the overproduction of alginate, a capsule-like polysaccharide (33,34), we measured the alginate levels to assess whether this surface-associated trait was altered in the mutant. The mutant *mreD^down^* did not produce alginate, resulting in a nonmucoid phenotype (Figure S3A-B). To optimize energy consumption, bacteria frequently acquire suppressor mutations under laboratory conditions, as previously described for alginate-associated sigma factor *algU* and its anti-sigma factor *mucA* (35,36). Whole-genome sequencing of *mreD^down^* revealed no mutations in alginate biosynthesis genes but identified a point mutation in the gene encoding ClpX, a component of the ClpXP protease complex that degrades MucA (Figure S3C) (37). In the wild-type strain, MucA degradation releases AlgU from the inner membrane, activating alginate biosynthesis genes as part of AlgU regulon (37). In *mreD^down^*, this process is likely impaired, leading to the loss of alginate production. Supplying *clpX* in *trans* restored alginate production in *mreD^down^* (Figure S3B), suggesting that the secondary mutation in *clpX* alleviated the energetic cost of alginate production while *mreD^down^* copes with a defective elongasome.

Overall, while maintaining cytotoxicity, *mreD^down^* exhibits impaired motility and loss of alginate production.

### Divisome overactivation compensates for the elongasome defect in *mreD^down^*

Previous studies reported that elongasome-targeting mutations or antibiotic treatments could be bypassed by the overexpression of division-associated genes. For example, *E. coli* became resistant to PBP2-targeting antibiotic, Mecillinam, upon overproduction of FtsZ septal division protein (38,39). Similarly, the overexpression of *ftsQAZ* genes in *E. coli* suppressed the lethality of deletions in the *mre* operon (22).

We wondered whether the *mreD^down^* strain withstands the downregulation of the essential *mreD* gene through overactivation of the divisome. To test this hypothesis, we analyzed the ability of the mutant to grow in the presence of Aztreonam, an antibiotic targeting PBP3, the divisome bPBP (Figure 2A and 2B). Wild-type *P. aeruginosa* strain tolerated a 4-hour incubation with Aztreonam at 5 µg/mL, and became significantly elongated, compared to untreated bacteria, as expected in the case of impaired division (Figure 2A and 2B). On the other hand, *mreD^down^* was significantly more sensitive to Aztreonam, exhibiting a four-log decrease in colony forming units (CFU) in presence of the antibiotic. *mreD^down^* also displayed pronounced morphological alterations, including irregular cell surfaces and multiple membrane swellings distributed along the length of the cell (Figure 2A and 2B). In comparison, both strains tolerated equally A22, an antibiotic targeting MreB (Figure 2A). This treatment caused the formation of spherical cells in the case of the parental strain and no change in morphology for *mreD^down^* (Figure 2B). These results show that *mreD^down^* depends primarily on its divisome to survive.

**Figure 2:**
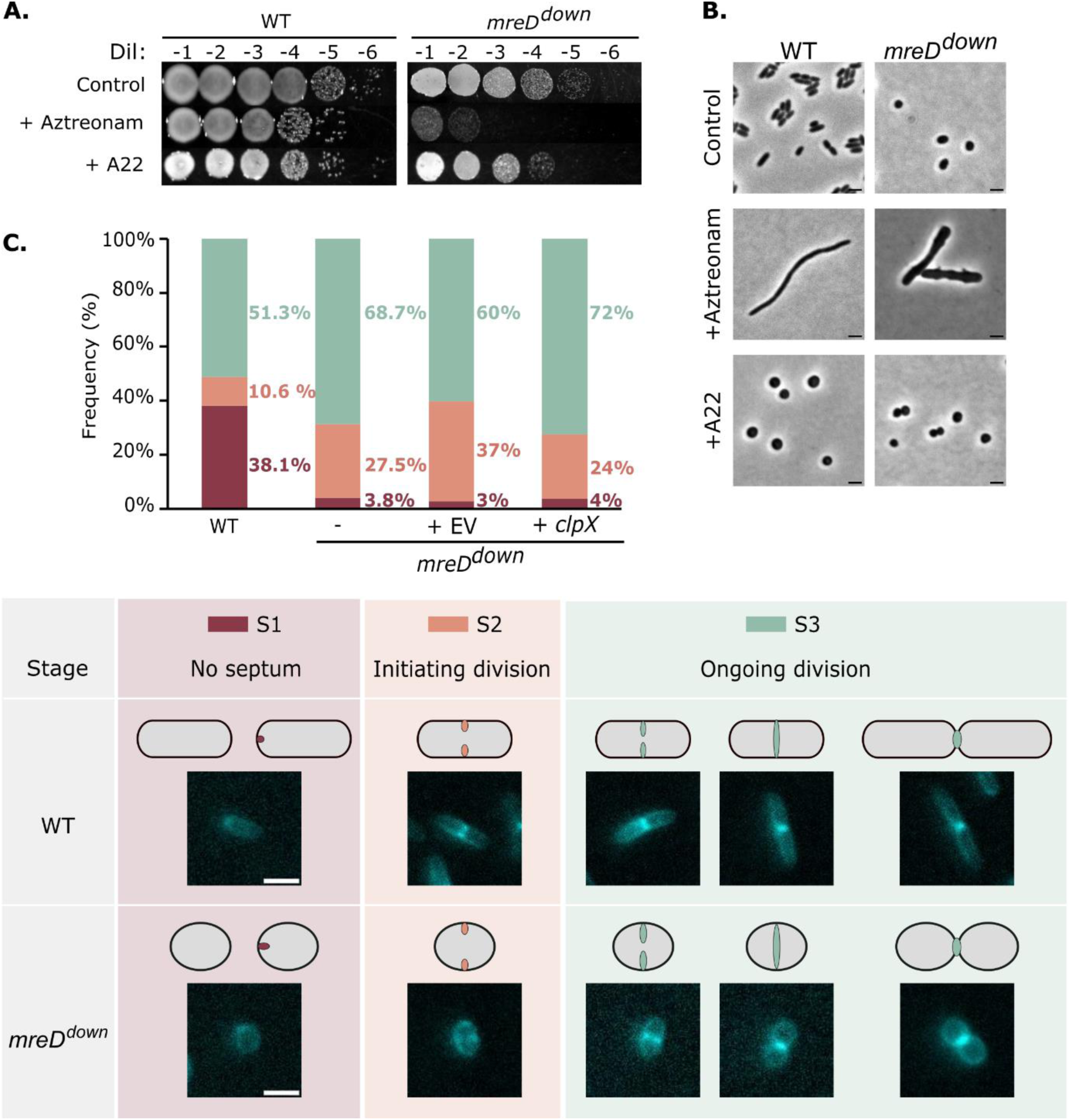
m*r*eDdown presents an overactivated divisome. **A.** Wild-type (WT) and *mreD^down^* were treated with 5 µg/mL Aztreonam (targeting the divisome) and 125 µg/mL A22 (targeting the elongasome). After 4 hours of incubation, cultures were serially diluted and spotted onto LB agar. Dil: dilution. **B.** Phase-contrast microscopy of the treated bacterial cells after 4 hours of incubation with the indicated drugs. Scale bar: 2 µm. **C.** Quantification of bacterial division stages based on HADA labeling: no septum (S1), initiating division (S2), and division in progress (S3) (n= 1513 (WT), 1067 (*mreD^down^*), 953 (*mreD^down^*/pSW-EV), and 629 (*mreD^down^*/pSW-*clp*X)). Scale bar: 2 µm.

Previous studies in *E. coli* described that targeting PBP2 with Mecillinam, not only results in the impairment of the elongasome activity but also triggers a futile cycle in the PG metabolism (40). This causes continuous degradation of cell wall material and depletion of PG precursors required for growth (40,41). To determine whether Mecillinam induces comparable effects in our system, we assessed the sensitivity of wild-type and mutant strains to this antibiotic (Figure S4). Both strains resisted to 450 µg/mL of Mecillinam. This resistance may reflect known features of this species, such as β-lactamase activity, and the limited efficacy of Mecillinam against *P. aeruginosa* (42,43). Overall, Mecillinam-induced elongasome inhibition in *E. coli* does not appear to elicit a comparable response in *P. aeruginosa*.

Next, we quantified the number of bacteria across three stages (S) of growth: non-dividing cells lacking a septum (S1), cells initiating division (S2), and cells undergoing an active division (S3) (Figure 2C). To do that, new septal PG synthesis was monitored through the incorporation of the fluorescently labeled D-amino acid, 7-hydroxycoumarin-3-carboxylic acid-amino-D-alanine (HADA) (44). In the parental population, 38.1% of cells were classified in S1, showing no septum at midcell, while only 10.6% of cells had initiated septum formation (S2). The remaining 51.3% of wild-type cells were in advanced division stages with extended septa (S3). In contrast, we counted ten-fold less non-dividing *mreD^down^* cells (S1). Instead, 27.5% were in S2 and 68.5% were actively dividing (S3) (Figure 2C and Figure S5). Interestingly in the mutant, a second division cycle was occasionally initiated prior to the completion of the first cell cycle (Figure S6). The increased frequency of dividing cells indicated enhanced divisome activity, consistent with the reliance of the mutant on this PG synthesis machinery for growth.

Since previous studies have shown that FtsZ interacts with and is negatively regulated by ClpXP protease complex (45–47), we also monitored PG synthesis using HADA incorporation in *mreD^down^* supplemented with *clpX in trans.* The distribution of cells through the different growth stages was similar to that of the *mreD^down^* mutant, with only 4% of cells in the non-dividing stage (S1), 24% initiating division (S2), and 72% in advanced stages of division (S3) (Figure 2C). These results indicate that divisome overactivation in *mreD^down^* occurs independently of ClpX expression.

Considering that *mreD^down^* displays an overactivated divisome and a defective elongasome, we wondered whether these alterations would affect the structural properties of the PG layer in the mutant. We therefore purified PG sacculi of the two strains at exponential phase, using boiling SDS and protease treatment, as previously described by Corona and Vollmer (48). Isolated sacculi thickness was measured using Atomic Force Microscopy (AFM) (Figure 3). Sacculi from parental cells displayed an average height of 8 nm. In contrast, *mreD^down^* sacculi showed a significantly reduced thickness (∼4 nm) (Figure 3A). These results indicate that, despite the increased divisome activity, *mreD^down^* is incapable of incorporating the adequate amount of lipid II in the periplasm, resulting in a thinner cell wall.

**Figure 3:**
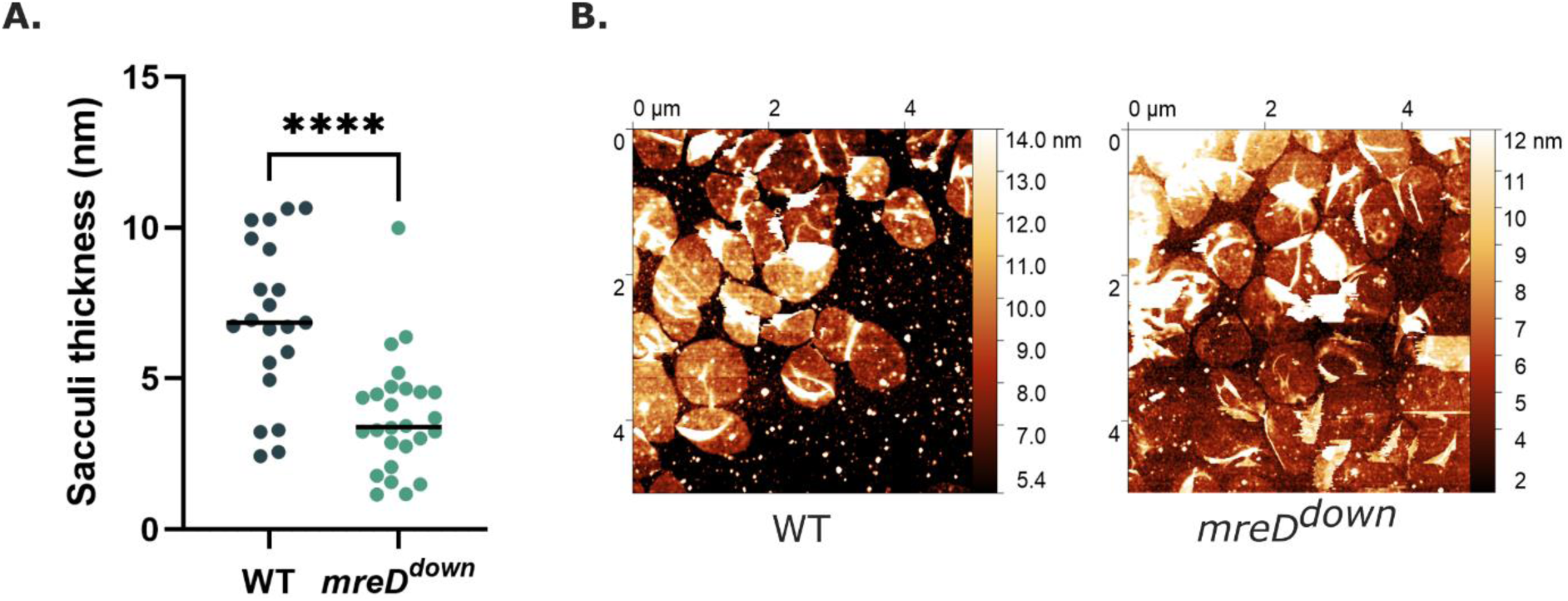
m*r*eDdown displays a thinner peptidoglycan layer. **A.** Quantification of sacculi thickness measured by atomic force microscopy (AFM). Each point represents an individual sacculus, and horizontal lines indicate the mean value. Statistical significance was determined using an unpaired t-test; ****p<0.0001. **B.** Representative AFM topography images of isolated sacculi from exponential-phase cultures of the WT and *mreD^down^* strains. AFM measurements were conducted on at least n= 25 sacculi per strain, using two independent preparations.

### Synthetic lethality screen identifies multiple genetic interactions between PG synthesis genes

To investigate *P. aeruginosa* responses and coping mechanisms in the presence of a defective elongasome, we performed a synthetic lethality screen in the *mreD^down^* background. Transposon mutant libraries were generated using a Himar-1 transposon in both the wild-type and *mreD^down^* strains, followed by Tn-seq analysis to assess mutant fitness. In the *mreD^down^* background, each clone carries a transposon (Tn5-Tc) insertion in *mreC* along with a second transposon (Himar-1-Gm) inserted elsewhere in the genome. The fitness of these double transposon mutants is then compared to that of single transposon mutants in the wild-type background. Beneficial and detrimental genetic interactions across the *P. aeruginosa* genome are depicted on the chord plot (Figure 4A, Table S2). Green links indicate double mutants with increased fitness, whereas orange links represent detrimental or synthetically lethal mutations. Validating the screen, transposon insertions in *ponA*, encoding PBP1a, produced the strongest fitness benefit in *mreD^down^* (Log_2_ (fold change (FC)) = 7.6), while insertions in *ponB* (encoding PBP1b) were highly detrimental (Log_2_ (FC) = −4.5) (Figure 4B-C, Table S2). As previously described, PBP1a is an aPBP that interacts with and stimulates components of the elongasome, including PBP2 (13). In *Bacillus subtilis*, mislocalization of the PBP1a homolog, PBP1 contributes to the shape defects and membrane bulges observed in an *mreB* deletion mutant (23). PBP1a and PBP1b are partially redundant and synthetically lethal (49). Thus, mislocalization of PBP1a in *mreD^down^* likely renders mutations in *ponB* highly detrimental, consistent with our data (Figure 4B-C, Table S3).

**Figure 4.**
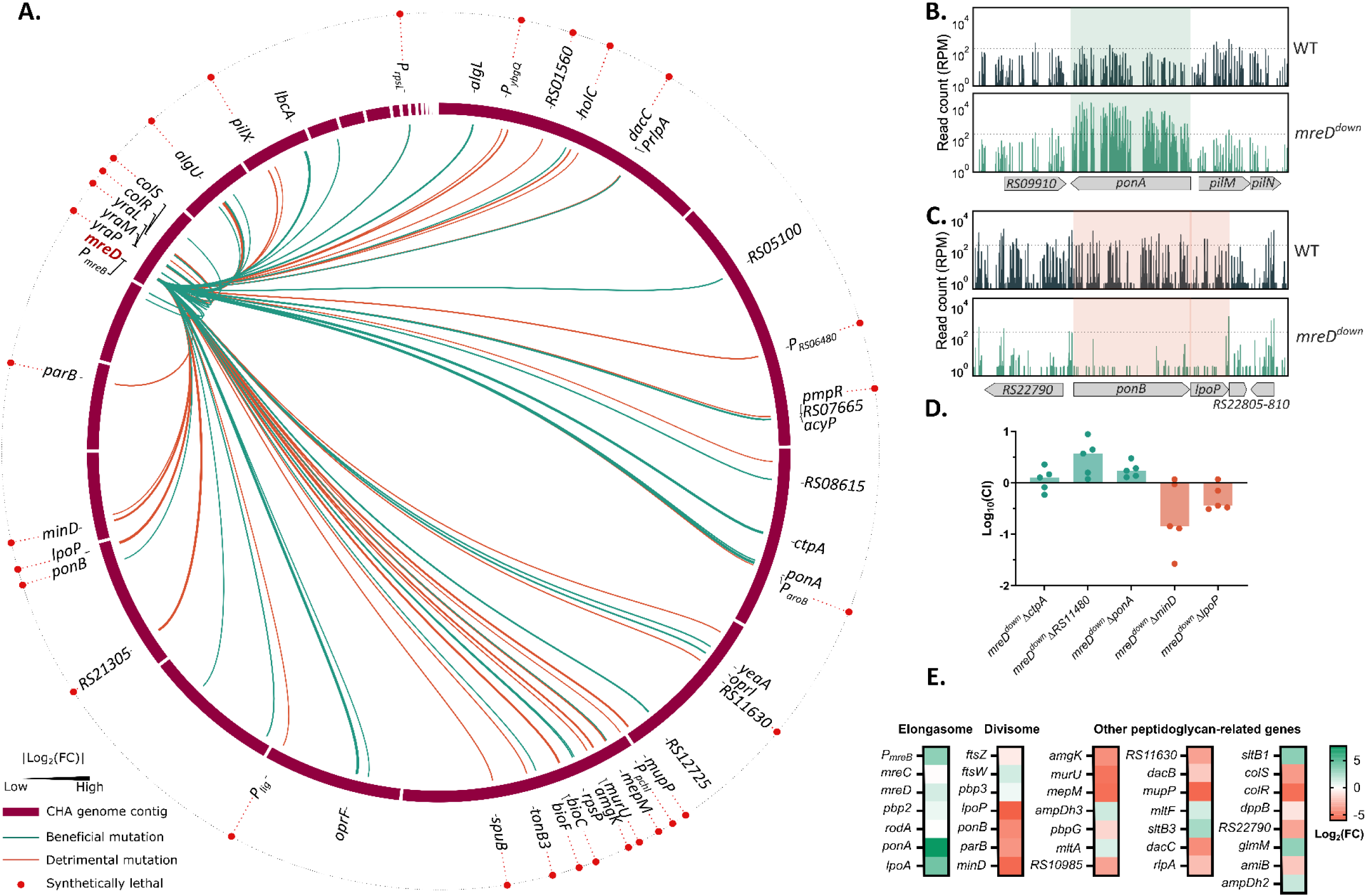
Synthetic lethality screen in an elongasome defective mutant *mreD^down^.* **A.** Chord plot representation of the synthetic lethality screen. Performed with Circos on usegalaxy.eu. **B.** Tn-seq profile of genomic region surrounding the *ponA* gene, showing the number of normalized reads in wild-type (WT) and *mreD^down^*. **C.** Tn-seq profile of genomic region surrounding the *ponB* and *lpoP* genes, showing the number of normalized reads in WT and *mreD^down^*. **D.** Competition index of clean deletion mutants *mreD^down^*Δ*ctpA*, *mreD^down^*Δ*RS11480, mreD^down^*Δ*ponA, mreD^down^*Δ*minD* and *mreD^down^*Δ*lpoP* in competition with the parental *mreD^down^* strain. Each dot represents a biological replicate, and the histogram marks the median. **E.** Heatmap showing Tn-seq results of the *mreD^down^* vs the WT strain (Log_2_(FC) values) for peptidoglycan (PG)-related genes. For elongasome and divisome, all values were used (no selection filter). For ‘Other PG-related genes’, only hit with |Log_2_(FC)|>1 and *P_adj_*<0.05 were used.

Similarly, *lpoP*, encoding the outer membrane lipoprotein that activates PBP1b, was synthetically lethal in the *mreD^down^* background (Figure 4A and 4C, Table S3) (50,51). Additionally, transposon insertion in the promoter region of the *mreBCD* operon, likely leading to operon overexpression due to the internal promoter within the transposon, resulted in increased fitness of *mreD^down^* (Log_2_ (FC) = 3.47, Table S2).

To further validate the screen, we selected three beneficial secondary mutations (*ponA*, *ctpA*, and the gene coding for putative L, D-transpeptidase *RS11480*/*PA2854*) and two detrimental ones (*lpoP* and *minD*) and assessed their fitness. In competition assays with *mreD^down^*, all engineered mutants behaved as predicted from the Tn-seq results, further validating the approach for investigating *P. aeruginosa* responses to a defective elongasome (Figure 4D).

### Identification of key pathways in an elongasome-defective mutant

Functional enrichment analysis of the most significant hits (|Log_2_FC| > 1; *P_adj_* < 0.05) using the STRING database revealed a strong enrichment of PG-related metabolic processes among both beneficial and detrimental interactions (Figure S7A) (52). Additionally, the data suggested that disruption of genes coding for quinone-binding and NADH dehydrogenase proteins increased the fitness of *mreD^down^* (*nuoN*: Log_2_(FC) = 3.44; *nuoG*: Log_2_(FC) = 3.07; Figure S7B, Table S2), whereas disruption of genes involved in DNA repair, stress response, and nucleic acid metabolic processes was detrimental (*recG*: Log_2_(FC) = -2.28; *uvrC*: Log_2_(FC) = -1.29; Figure S7C, Table S2). In line with these findings, perturbation of elongasome-mediated PG synthesis compromises cell wall integrity and activates envelope stress responses (53). Envelope stress responses are tightly linked to cellular redox homeostasis, notably through the regulation of respiratory complexes such as NADH dehydrogenases, which are major sources of reactive oxygen species (ROS) (54,55). This relationship may explain the beneficial effect of disrupting *nuo* genes, potentially through reduced ROS production. Interestingly, both genes of the *colRS* regulatory two-component system, which detects and responds to envelope perturbations including antimicrobial stress, were identified as synthetic lethal hits in the screen (Figure 4A, Table S3) (56). Because ROS can induce DNA damage, bacteria rely on DNA repair pathways to maintain genomic integrity under stress conditions. Accordingly, disruption of DNA repair genes was detrimental in the *mreD^down^* background (55).

Genes encoding endopeptidases and hydrolases, and the dipeptide transporter system, were also enriched among detrimental hits. Notably, *dacC,* encoding the low-molecular-weight PBP6 carboxypeptidase, was synthetically lethal in *mreD^down^* (Figure 4A, Table S3). In addition, disruption of all four genes encoding the dipeptide permease (Dpp; *dppBCDF*) significantly decreased fitness (Table S2). The Dpp system mediates the transport and utilization of di- and tripeptides, an important step upstream of PG recycling, which may become particularly critical in the *mreD^down^* context (57). Disruption of several hydrolases also impaired fitness, further underscoring the importance of PG turnover.

Given the strong enrichment of PG metabolic processes in the synthetic lethality screen, we focused subsequent analyses on PG-related pathways. As shown in Figure 4E, disruption of genes involved in multiple aspects of PG homeostasis, including elongasome genes, divisome genes and other PG-related genes, significantly impacted fitness.

Overall, *mreD^down^* cells appear to rely on stress response and DNA repair pathways to cope with the consequences of a defective elongasome. More importantly, our analysis identifies conditionally essential genes involved in PG synthesis and recycling that are critical for maintaining cell envelope integrity under these conditions.

### MepM becomes essential upon elongasome dysfunction

Among the synthetically essential genes, this screen identified *mepM* (*RS13525/PA0667*) (Figure 4A, 4E, and 5B, Table S3), which encodes a cell wall hydrolase. MepM is a D,D-endopeptidase that cleaves peptide crosslinks in PG (58) (Figure 5A.1). MepM activity is tightly regulated by the CtpA/LbcA system, a homologue of the well characterized Prc/NlpI system in *E. coli,* where LbcA activates CtpA, promoting MepM degradation (Figure 5A.2 and 3) (58–61). In agreement, our screen revealed an enrichment of mutants with transposon insertions in *ctpA* and *lbcA* genes upon elongasome defects (Figure 5B). These results show that preventing MepM degradation, thereby maintaining its physiological levels, benefits the *mreD^down^* mutant. This increase in fitness occurs despite MepM-like endopeptidases being functionally linked to the lateral wall synthesis (62–64). In *E. coli*, the endopeptidase activity of MepS has been shown to antagonize cell division while promoting cell elongation by cleaving peptide crosslinks within the PG mesh, thereby creating space for the insertion of new strands during lateral wall expansion (62). Consistent with this model, production of a single *E. coli* endopeptidase, out of the eight encoded in the genome, reduced lateral wall expansion, whereas deletion of all eight enzymes resulted in spherical morphology (63). Together, these findings indicate that the endopeptidase activity is required for lateral wall synthesis and maintenance of rod-shaped morphology.

**Figure 5:**
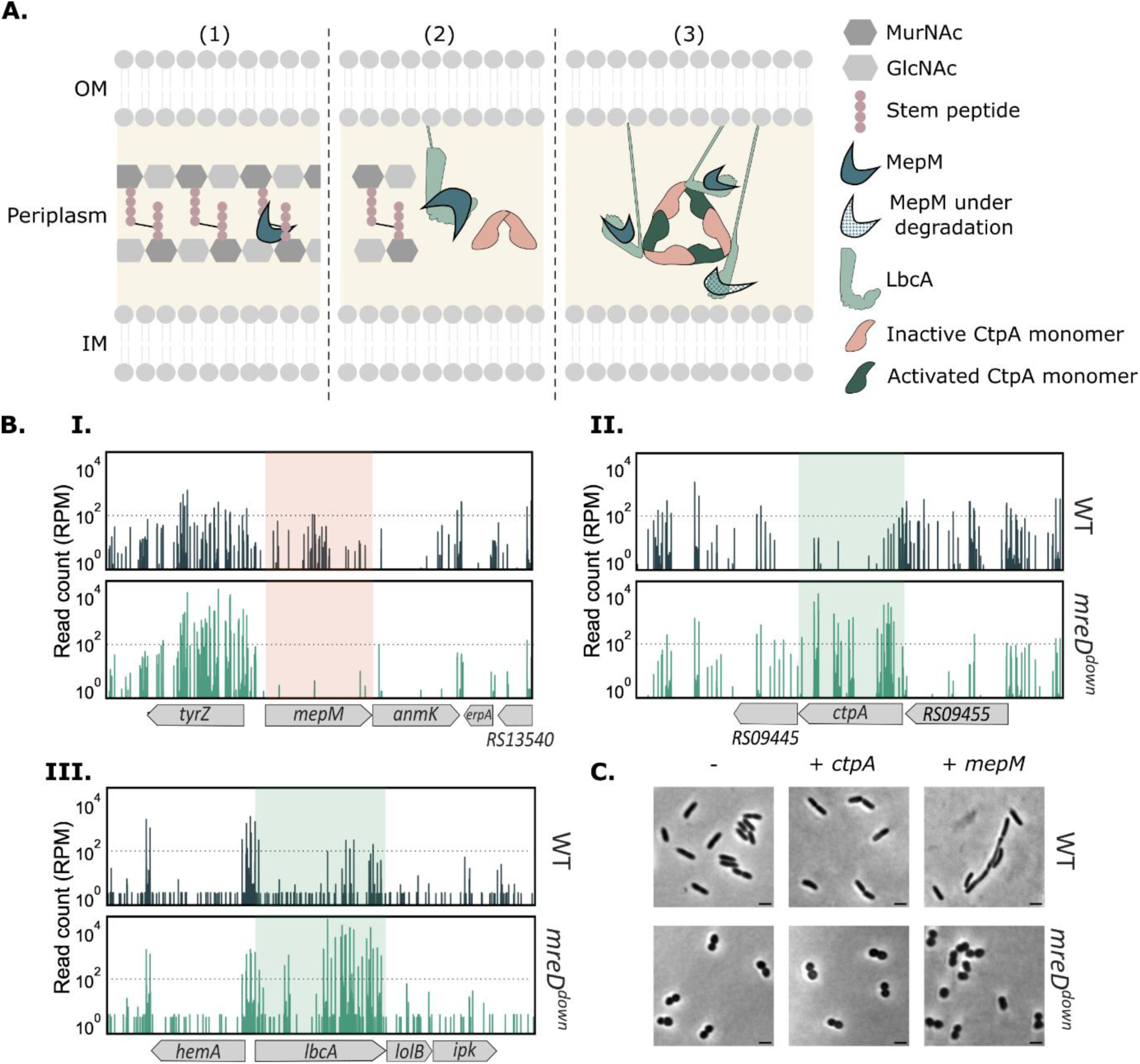
m*e*pM is essential in the elongasome-impaired *mreD^down^* mutant. A. Schematic model of CtpA activation by LbcA leading to MepM degradation proposed by (61). (1) MepM is a periplasmic endopeptidase that cleaves peptide crosslinks within the peptidoglycan. (2) MepM associates with membrane-anchored LbcA prior to CtpA binding. CtpA is present in the periplasm as an inactive dimer. (3) Upon interaction with a MepM-LbcA complex, three LbcA-bound CtpA dimers assemble into a triangular hexameric structure. Within each dimer, only one CtpA becomes activated (dark green), enabling proteolytic degradation of MepM. IM, inner membrane. OM, outer membrane. B. Tn-Seq profiles of genomic region surrounding the *mepM* (I), *ctpA* (II), and *lbcA* (III) genes, showing the number of normalized reads in WT and *mreD^down^*. Orange and green backgrounds stand for detrimental and beneficial mutations, respectively. C. Phase-contrast microscopy images of WT and *mreD^down^* strains overexpressing *ctpA* or *mepM* from an arabinose-inducible promoter. Overexpression was triggered with 0.5% arabinose for 2h prior to imaging. Scale bar: 2 µm.

These observations raise the question of whether MepM functions as an elongasome-associated endopeptidase in *P. aeruginosa* and why it becomes essential in a mutant in which elongation is impaired. To assess whether MepM promotes elongation in *P. aeruginosa*, *mepM* was overexpressed *in trans* in both the parental and mutant strains. This led to pronounced elongation in the parental strain, while no morphological changes were observed in the *mreD^down^* background. In contrast, *ctpA* overexpression in both parental and *mreD^down^* mutant strains did not lead to morphological changes. These results indicate that elevated levels of MepM in the periplasm promote lateral wall expansion and cell elongation in the parental strain but not in *mreD^down^* mutant.

### *mreD^down^* mutant relies on MepM-mediated PG recycling pathway

MepM role in promoting cell wall elongation while remaining essential in an elongasome-impaired mutant, prompted us to further examine MepM function in PG recycling, as it contributes to the generation of recyclable PG fragments (65). We therefore hypothesize that MepM is crucial for *mreD^down^* as it supplies an essential source of cell wall material via PG recycling.

As previously described, the PG layer is a highly dynamic structure that undergoes continuous turnover during bacterial growth (66). This remodeling starts by the endopeptidases and lytic transglycosylases (66,67) (Figure 6A), which release soluble muropeptide fragments, primarily composed of GlcNAc-1,6-anhydroMurNAc peptides (66). These fragments are then imported into the cytoplasm to be recycled (68). Recycling of sugar moieties occurs through two different routes in *P. aeruginosa*.

**Figure 6.**
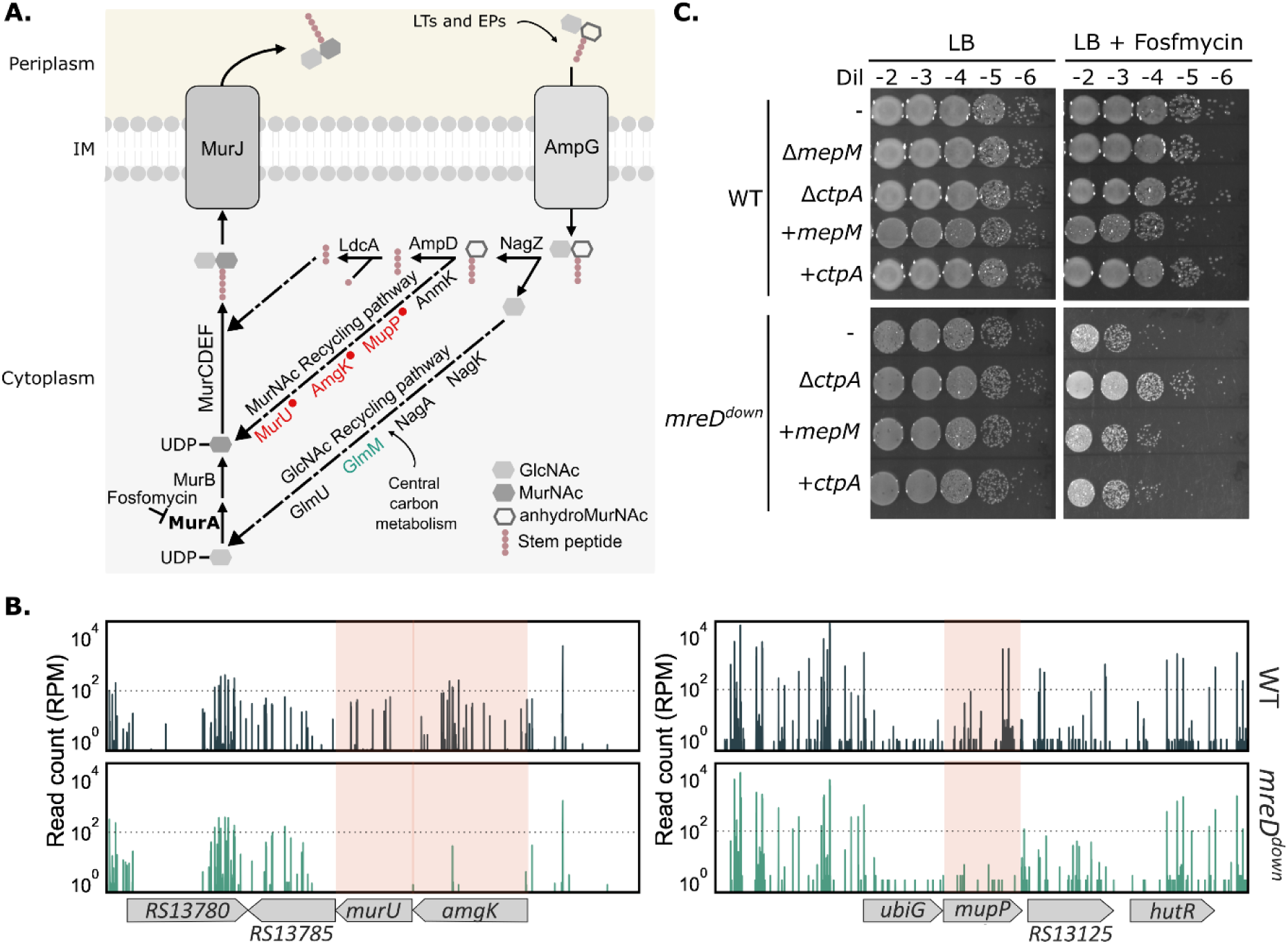
*mreD^down^* depends on an essential MepM-linked peptidoglycan recycling pathway. **A**. Scheme of the PG recycling pathway. During cell wall remodeling, PG cleavage by endopeptidases and lytic transglycosylases releases soluble muropeptides containing GlcNAc-1,6-anhydroMurNAc. These fragments are transported into the cytoplasm through the permease AmpG and processed by recycling enzymes. GlcNAc-derived metabolites are converted to UDP-GlcNAc through the Glm pathway, whereas 1,6-anhydroMurNAc is directly recycled into UDP-MurNAc through the MurNAc recycling pathway. Both routes replenish the cytoplasmic pool of peptidoglycan precursors used for cell wall biosynthesis. Genes shown in red with a red dot correspond to synthetic lethal mutations identified by Tn-seq, whereas the gene in green indicates a beneficial mutation. IM, inner membrane; LT, Lytic Transglycosylase, EP, Endopeptidase. The target of Fosfomycin, MurA is indicated in bold. **B.** Tn-seq profiles of the *murU*, *amgK*, and *mupP* loci in *Pa* WT and *mreD^down^* strains. Vertical lines above each locus represent individual transposon insertion sites, and line height corresponds to the number of sequencing reads recovered at each site. Orange backgrounds stand for detrimental mutations **C**. Fosfomycin sensitivity assay. *Pa* WT, *mreD^down^*, their corresponding deletion mutants (Δ*mepM* and Δ*ctpA*), and strains overexpressing *mepM* or *ctpA* were treated with 10 µg/mL Fosfomycin. After 4 h of incubation, cultures were serially diluted and 10 µL of each dilution were spotted onto LB agar plates. Dil: dilution.

GlcNAc is converted into Uridine Diphosphate (UDP)-GlcNAc (69–73) (Figure 6A), which is then funneled into the *de novo* biosynthesis pathway, where it can be used for the production of UDP-MurNAC (69). However, 1,6-anhydroMurNAc follows a more direct route in *P. aeruginosa* where it is converted directly to UDP-MurNAc through the MurU pathway (74–77).

The Tn-seq data revealed that *murU, amgK* and *mupP*, critical for the 1,6-anhydroMurNAc recycling, are synthetically lethal in an *mreD^down^* background (Figure 6B, Table S3). This indicates that the MurNAc PG recycling pathway, bypassing *de novo* PG synthesis, is essential in the absence of a fully functional elongasome. To test this hypothesis further, the wild-type strain, *mreD^down^*, their corresponding mutants Δ*ctpA* and Δ*mepM*, and the strains overexpressing *ctpA* and *mepM* were treated with Fosfomycin. The Δ*mepM* mutant in the *mreD^down^* background could not be generated, given *mepM* essentiality in this context, and was therefore not tested.

Fosfomycin blocks MurA, an enzyme responsible for the production of UDP-MurNAc precursor, forcing the bacteria to bypass the *de novo* PG synthesis through the MurU pathway. Under these conditions, the parental strain and its mutants had a moderate sensitivity to 10 µg/mL of Fosfomycin after a four-hour incubation (Figure 6C), whereas *mreD^down^* cells were significantly more sensitive. Interestingly, Δ*ctpA* mutation restored wild-type-like sensitivity to Fosfomycin in an *mreD^down^* background. These results indicate that preventing degradation of MepM during growth confers resistance to Fosfomycin in *mreD^down^* cells. On the other hand, the overexpression of *mepM* did not result in a similar phenotype, a pattern also observed in the parental strain, as it resulted in an increased sensitivity (Figure 6C). No differences were observed upon *ctpA* overexpression in either strain. These results suggest that PG recycling pathway is essential upon elongasome dysfunction and relies on physiological levels of MepM probably to supply the highly active divisome with the adequate amount of PG precursors.

## Discussion

*P. aeruginosa* causes life-threatening infections and resists β-lactams that target PG synthesis, highlighting the importance of studying this process (1). While the organization of the lateral PG synthesis machinery, the elongasome, is well characterized, how bacteria adapt to its alteration remains poorly understood. In this study, we show that the loss of a fully functional elongasome leads to divisome overactivation as a compensatory mechanism, in agreement with previous studies (22,38,39). We also show that MepM overexpression promotes lateral wall expansion in *P. aeruginosa* model, while maintaining physiological level confers resistance to Fosfomycin in an elongasome-impaired mutant. Together, these results propose an essential role of this endopeptidase in regulating adaptation to the loss of the elongasome, by favoring the MurU-PG recycling process over the *de novo* synthesis pathway.

The viable *mreD^down^* mutant provides a unique model to investigate the competitive interplay between the divisome and the elongasome. Cell growth relies on a dynamic balance between elongation and division, where disrupting one process actively drives the other. Here, we show that elongasome dysfunction in *P. aeruginosa* triggers a compensatory hyperactivation of cell division, and our Tn-seq results confirm conditional essentiality of division genes (Figure 2). This compensatory mechanism aligns with previous observations in *E. coli*, where a strain harboring a temperature-sensitive allele of *ftsI* elongate faster than a wild-type at permissive temperatures (62,78).

In addition to hyperactivation of the divisome, the endopeptidase-encoding gene *mepM* was also essential in *mreD^down^* background (Table S3; Figure 5B). MepM belongs to the M23 family of Zn²⁺-dependent metallopeptidases (79,80). BLAST analysis identified five homologues in *P. aeruginosa* (RS27260, RS01785, RS00905, RS10560, RS25165) sharing 32-40% identity with MepM as presented in Table S4. Among these, RS27260 and RS01785 retain the catalytic sequence motif (H(x)_n_D and HxH) whereas others show variations (79). In fact, M23 family members which lack the full set of catalytic residues, do not function as endopeptidases, but instead mediate protein activation through direct interactions with their substrates, as described for *E. coli* division proteins EnvC and NlpD (81,82). Tn-seq results indicate that transposon insertions in RS27260 and RS01785 genes are also detrimental, emphasizing the importance of MepM-like activities in *mreD^down^* (Table S4).

In the periplasm, MepM turnover is mediated by CtpA, which is activated by LbcA (Figure 5A) (58–61). Tn-seq analysis showed that insertions in *ctpA* and *lbcA* benefit the *mreD^down^* mutant, supporting the need to maintain physiological MepM levels for survival, likely by preventing its degradation by CtpA (Figure 5B).

Building on these findings, MepM-dependent regulation can thus be placed within the broader context of the proposed competition between the elongasome and the divisome. In the current model, competition is mediated by the shared lipid II pool, allowing one machinery to utilize the entire pool when the other is impaired (62). The pool of lipid II is sustained by both *de novo* PG synthesis and PG recycling (9). By showing that genes coding for key enzymes of the pathway (*murU, amgK*, and *mupP*) are synthetically lethal in the *mreD^down^* background (Table S3), our work establishes the MurNAc PG recycling route as a critical determinant when elongasome function is impaired and septal activity is elevated. Consistently, loss of CtpA (Δ*ctpA*) in the *mreD^down^* background confers resistance to Fosfomycin, an antibiotic that enforces dependence on PG recycling, thereby functionally linking MepM activity to the essential MurU-dependent pathway. These findings are further supported by recent work demonstrating that the disruption of PG recycling of *Caulobacter crescentus*, which possesses the MurNAc-PG recycling enzymes, reduces precursor availability and impairs cell division (9). However, whether PG recycling drives increased division by replenishing the lipid II pool, or whether it becomes essential as a consequence of divisome overactivation, remains an open question.

Further supporting this model, treatment of *E. coli* with Mecillinam, which targets PBP2 and thereby impairs the elongasome, triggers a futile cycle in PG metabolism. In this context, PBP2 inhibition leads to continuous synthesis of uncrosslinked PG that is rapidly degraded by lytic transglycosylases (40,41). The resulting fragments must re-enter the PG precursor pool via the multi-step *de novo* pathway to be regenerated. However, this cycle cannot match the rapid rate of PG degradation, ultimately impairing cell wall synthesis and causing cell death due to the depletion of precursors (40). Importantly, increasing endopeptidase activity, through MepS overexpression, confers Mecillinam resistance, demonstrating that modulation of endopeptidase activity can influence bacterial survival under elongasome impairment (41). In contrast, our results, in *P. aeruginosa,* reveal that endopeptidase function is required at physiological levels, rather than through overexpression, to sustain growth under similar conditions. Unlike *E. coli*, where precursor depletion is lethal, *P. aeruginosa* likely mitigates this imbalance through the MurU-dependent recycling pathway, enabling more efficient replenishment of PG precursors. In this context, MepM likely promotes the generation of recyclable muropeptides, thereby sustaining precursor pools and supporting survival.

In addition, our data identify the divisome-associated aPBP PonB and its activator LpoP as major determinants of fitness in the *mreD^down^* background. Consistently, Lai *et al.* further demonstrated that increased endopeptidase activity enhances cell wall synthesis by promoting aPBP-mediated PG insertion (41). Together, these results support a model in which endopeptidase activity functionally cooperates with aPBPs to sustain peptidoglycan synthesis under elongasome-defective conditions.

Overall, PG precursor production, in both the periplasm and the cytoplasm, appears to be tightly regulated and plays a central role in adaptation to stress. Under wild-type conditions, PG recycling and *de novo* PG synthesis jointly supply both elongasome and divisome with sufficient PG units (Figure 7A). When *mepM* is overexpressed, elongation is favored, leading to excessive bacterial elongation (Figures 7B), although the preferred precursor source remains unclear. In contrast, when the elongasome is impaired, as in *mreD^down^* mutant, the divisome becomes dominant to compensate for the impaired elongasome machinery. In this context, the MepM-dependent PG recycling pathway is favored over the *de novo* PG synthesis (Figure 7C). This may reflect differences in the rate at which lipid II building blocks are produced by the two processes, or even differences in the energy required to sustain them. These are mechanistic details that remain to be determined.

**Figure 7:**
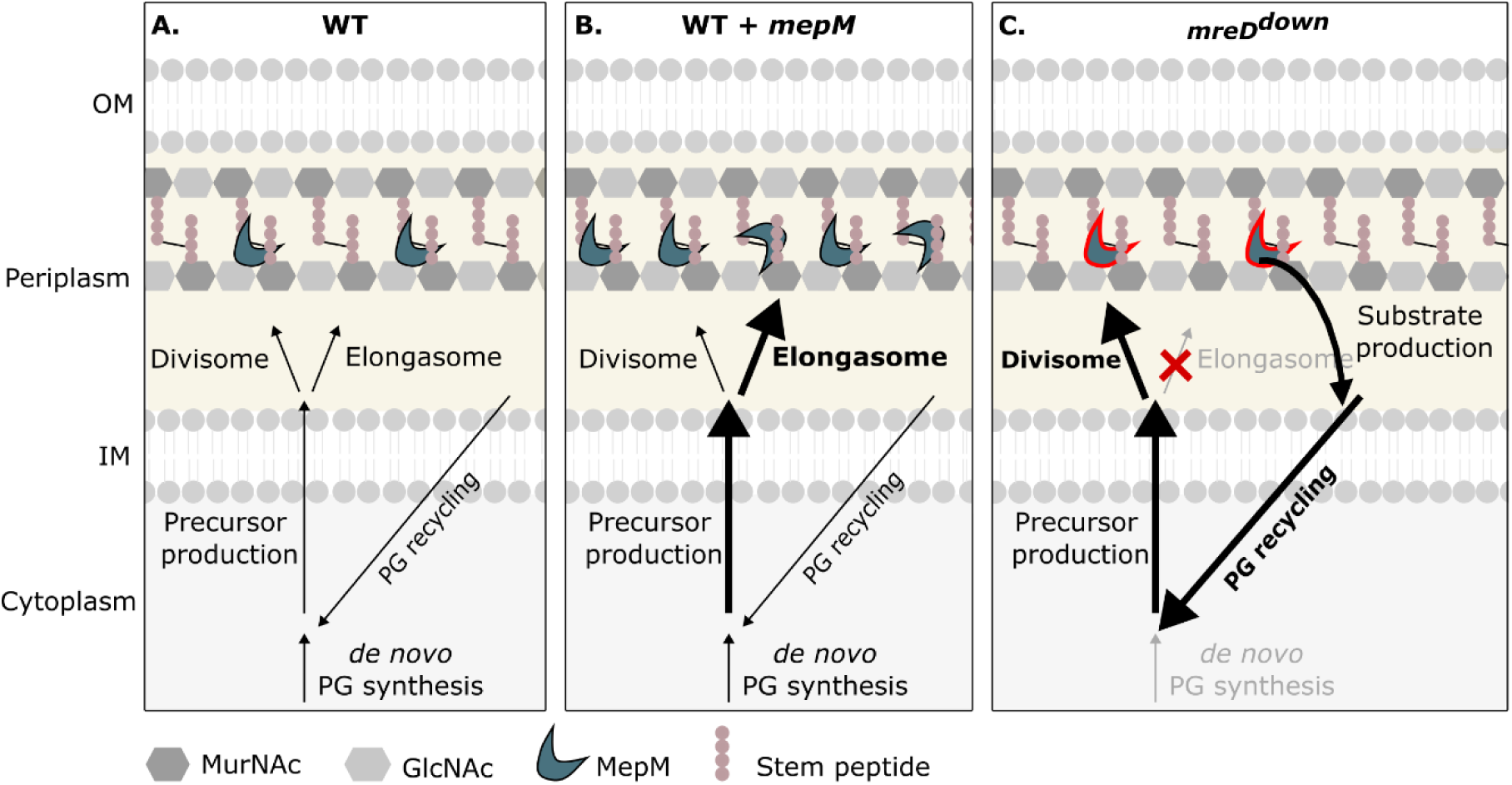
MepM-dependent PG recycling sustains divisome when elongation is impaired in *Pseudomonas aeruginosa*. **A.** In wild-type (WT) cells, elongasome and divisome activities are balanced. PG precursors are produced via *de novo* synthesis in the cytoplasm and supplemented by PG recycling, providing sufficient material for both elongation and division. **B.** Overexpression of *mepM* favors elongasome activity, enhancing lateral wall expansion. The demand for PG precursors increases, but both *de novo* synthesis and recycling pathways remain active to support growth. **C.** Downregulation of *mreD* impairs elongasome function, leading to overactivation of the divisome. PG recycling, dependent on MepM, becomes the dominant source of precursors to sustain septal synthesis, while *de novo* synthesis contributes to a lesser extent. The red outline of MepM indicates essentiality. OM: Outer membrane, IM: Inner membrane.

Previous studies have highlighted the importance of endopeptidases under stress conditions. For instance, enzymes such as ShyA of *Vibrio cholerae* can promote cell wall degradation upon antibiotic treatment (83), while others, as MepM and MepS of *E. coli*, become essential under various stress conditions such as salt and EDTA, respectively (84). Here, we extend this concept by showing that endopeptidase activity is also critical in a context where elongation is impaired. Our results identify MepM as a key adaptive determinant that maintains cell wall remodeling and peptidoglycan homeostasis despite elongasome dysfunction. This establishes a model in which endopeptidase-dependent recycling becomes essential to sustain growth when the balance between elongation and division is disrupted in *P. aeruginosa*.

Together, these findings reinforce the emerging view that endopeptidases and PG recycling enzymes act as versatile and adaptive regulators of cell wall homeostasis, true “jokers” of bacterial physiology, and highlight them as promising targets for the development of novel or combination antibacterial strategies.

## Material and methods

### Bacterial strains and growth

Bacterial strains and plasmids used in this study are listed in Table S5. Unless stated otherwise, bacteria were grown in Lysogeny Broth (LB) at 37°C with shaking. Antibiotics were used at the following concentrations: 75 µg/mL Gentamicin for *P. aeruginosa* and 50 µg/mL Gentamicin, 100 µg/mL Ampicillin for *E. coli*. 75 or 10 µg/ml Tetracycline for *P. aeruginosa* and *E. coli*, respectively. Irgasan-containing LB plates (25 µg/mL) were used to select for *P. aeruginosa*, when needed.

For growth curves, bacteria were grown overnight in LB at 37°C under agitation. Agarose (1%) was poured into an Edge 96-well plate (Nunc #056897), and bacteria were inoculated at an optical density at 600 nm (OD_600_) of 0.001, under indicated conditions. Plates were covered with breathable film (Breathe-easy, Diversified Biotech) and incubated for 20 hours in the spectrophotometer (Safas, MP96 Monaco) at 37°C under agitation (5 Hz). Bacterial growth was monitored by the absorbance at 600nm and the OD_600_ was measured every 10 minutes for 20 hours.

### Generation of the elongasome structural model

Figure 1A was generated using Pymol by integrating available structural data and predicted models. The PBP2/MreC interaction is represented using the active conformation [PDB 5lp5] (85), while the PBP2-TM/RodA and the MreC-TM/MreD interactions were modeled based on available structures [PDB 6pl5 and PDB 9dvb] (17,20), respectively. MreB oligomers were incorporated from published structure [PDB 1JCG] (14). Models of PBP1a (PA5045) of PAO1 and its full-length MreC were obtained using AlphaFold 3. The C-terminal proline-rich domain of MreC, predicted to be largely disordered, was omitted for clarity. Orientations of the periplasmic domains of MreC and PBP2 were manually adjusted based on the model proposed by Shlosman *et al.* (20).

### Genetic manipulations

The gene of *mreC* from *P. aeruginosa* CHA with native RBS was amplified by PCR with specific primers (Table S6) and cloned into *Eco*RI- and *Sac*I-digested pJN105 empty vector, giving pJN105-RBS-*mreC*. Then the RBS of *mreC* was replaced by the RBS of *tssK* from T6SS by mutagenesis (86), giving pJN105-RBS*_tss_*_K_-*mreC.* In addition, in the mutagenesis primers, an *Nde*I restriction site was added downstream the RBS*_tssK_* to allow the cloning of *mreD* and *mreCD* genes into the pJN105-RBS_tssK_ (Table S6). To generate the plasmid expressing *clpX, mepM-mCherry,* and *ctpA-mCherry,* their coding sequences were amplified by PCR (Tables S5 and S6). Their products were cloned into *SmaI*-digested pSW196 by sequence-and ligation-independent cloning (SLIC) (87). All constructions were verified by sequencing. pJN105- or pSW196- derived plasmids were introduced into *P. aeruginosa* by transformation or triparental conjugation, using pRK600 as a helper plasmid. To create deletion mutants, upstream and downstream flanking regions (approximately 500 bp) were amplified from genomic DNA by PCR using appropriate primer pairs (sF1/sR1, sF2/sR2 (Table S6)). Overlapping fragments were cloned into *Sma*I-digested pEXG2 by SLIC. Plasmids were used to transform *E. coli* competent cells, and their sequences were verified (Eurofins). Allelic exchange vectors derived from pEXG2 were introduced into *P. aeruginosa* by triparental mating, using pRK600 as a helper plasmid. Merodiploids, resulting from homologous recombination, were selected on LB agar plates containing irgasan and the appropriate antibiotic. Single merodiploid colonies were streaked on NaCl-free LB plates with 10% sucrose (w/v) to select for plasmid loss. Resulting sucrose-resistant clones were screened for antibiotic sensitivity and gene deletion by PCR.

### MreC expression

Overnight bacterial cultures were diluted into 30 mL of LB medium to an initial OD_600_ of 0.1 and grown at 37°C with shaking. Samples corresponding to the same number of bacterial cells (3 × 10⁷ CFU) were collected every 30 min starting from mid-exponential phase (OD_600_ = 1.0) up to stationary phase (OD_600_ > 3) and analyzed by immunoblotting. Proteins were separated on 12% SDS-PAGE gels, transferred to PVDF membranes, and blocked overnight in 5% (w/v) milk solution. Proteins of interest were labeled using targeted primary antibodies at the following dilutions: rabbit anti-MreC antibodies (Martins et al, 2021, doi: 10.1038/s41467-021-22957-9, 1:10,000), mouse anti-EF-Tu (Hycult Biotech, 1:10,000). Anti-rabbit-HRP (Sigma, 1:20,000) and anti-mouse-HRP (Sigma, 1:20,000) were used as secondary antibodies. Membranes were developed using the Luminata Classico Western HRP substrate (Millipore). The EF-Tu signal was used as a loading control.

### Study of bacterial morphology

#### Microscopy observations

Bacteria were grown in LB at 37°C with shaking, in presence of 0.3% arabinose, when indicated. In mid-exponential phase (OD_600_=1.0), bacteria were either observed directly or labeled with SYTO24 (1:5000 dilution from a 5 mM stock solution, ThermoFischer Scientific #S7559, Em 515nm) for 15 min in the dark at room temperature, then washed with phosphate-buffered saline (PBS, Gibco #14190-094). Bacteria were then mounted on Labteck (Thermo Fisher Scientific) under 1.5% low-melting agarose pads and visualized using an inverted X71 Olympus microscope controlled by the CellR Olympus system, automated in x, y, z axis and driven by Xcellence software (Olympus).

#### Size analysis

Image analysis was performed using ImageJ/Fiji (88) and quantitative bacterial morphology analysis was performed using the MicrobeJ plugin (28). Data on length and mean width were extracted, and an L/S ratio was calculated, reflecting cell roundness (Long/Short axis). For analysis of spherical bacteria, the parameters of maximum length were changed to allow correct segmentation. The number of bacteria was calculated from 6-10 independent images for each sample. At least 450 cells were analyzed in each sample.

#### RT-qPCR

Total RNA was extracted from 2 mL of exponential-phase bacterial cultures (OD_600_ = 1) using the TRIzol Plus RNA Purification Kit, followed by TURBO DNA-free treatment (Invitrogen). cDNA synthesis was performed using SuperScript IV Reverse Transcriptase (Invitrogen). Quantification was carried out by real-time PCR using SYBR Green chemistry with gene-specific primers (Table S6) and Luna Universal qPCR Master Mix (NEB). The absence of genomic DNA contamination was confirmed by including no-reverse transcriptase controls. Relative mRNA levels were determined using CFX Manager software (Bio-Rad) with *rpoD* as the reference gene.

#### Whole-genome sequencing

DNA libraries were prepared using the Collibri PS DNA library prep kit for Illumina according to the provider’s recommendation (Invitrogen). Briefly, DNA was mechanically sheared at 4°C by sonication (15 s pulses, 25 min total). Fragment size and concentration were assessed using high-sensitivity DNA chips on an Agilent Bioanalyzer 2100 (Agilent Technologies). Fragmented DNA was then end-repaired and dA-tailed, followed by ligation to dual-indexed adapters. Adapter-ligated fragments were purified and size-selected (150–800 bp) using DNA Cleanup Beads (Invitrogen). Libraries were amplified by PCR and purified again using DNA Cleanup Beads. Final library quality and size distribution were verified using high-sensitivity DNA chips on the Agilent Bioanalyzer 2100. Libraries were sequenced on an Illumina NextSeq 500 (I2BC, Saclay Paris). Sequencing was performed in paired-end reads (2x150nt) with a theoretical coverage depth of 100X per genome. The raw data has been deposited to the Sequence Read Archive (SRA) under the NCBI bioproject number PRJNA1445359. Sequencing reads were analyzed on the Galaxy platform (The Galaxy Community, 2022). Reads were trimmed using fastp (89); we used FastQC for read quality control (90); SPAdes for genome assembly (91); Quast for quality control of the assembled genome (92); Prokka for gene annotation (93), and Snippy for assessments of SNPs (94).

#### Motility assays

Bacteria were streaked out from single colonies and incubated with appropriate antibiotics. Swimming plates containing Tryptone 10 g/L (Difco), Agarose 0.3 % (Difco), NaCl 5 g/L (Sigma) were allowed to equilibrate at room temperature in the morning and bacteria were carefully inoculated at mid-height with a toothpick. Plates were sealed with parafilm and bacteria were allowed to swim at 30°C.

#### Immunofluorescence

For FliC immunofluorescence, bacteria were seeded on a coverslip in each well of a 24-well plate and grown statically for 2h at 37°C. Bacteria were fixed with 4% paraformaldehyde (PFA) for 48h at 4°C. Bacteria were then incubated with 2% BSA (in PBS) for 30 min, and further incubated with anti-FliC antibodies (1:250 dilution, (95)) for 45min stained using standard procedure with appropriate primary and secondary antibodies. Images were collected on a Zeiss Axioplan equipped with an objective x100 numerical aperture 1.3 oil immersion objective.

#### Cytotoxicity assay

The cytotoxic activity of strains was evaluated using macrophage cell line J774A.1 (ATCC, TIB-67). Cells were seeded in 96-well plates at 5x10⁴ cells per well and incubated overnight. The following day, the medium was replaced with fresh DMEM supplemented with propidium iodide (1:1,000 dilution of 1 mg/ml stock solution, SIGMA#P4864). Bacterial cultures grown to an OD_600_ of 1.0 were diluted to obtain a multiplicity of infection (MOI) of 5 and added to the cells. Cytotoxicity was monitored at 37°C and 5% CO_2_ after 2h 30min by Lactate dehydrogenase release into the supernatant using Cytotoxicity Detection Kit (Roche Applied Science). The 100% cell death value was quantified after addition of 1% Triton X-100 in control wells, in triplicate.

#### Alginate quantification

Saturated bacterial cultures were plated onto LB agar P100 plates. For strains harboring the pSW196 vector and derivatives, LB was supplemented with 0.5% (w/v) arabinose. Following incubation overnight at 37°C, bacterial biomass was scraped from the agar surface using a sterile loop and resuspended in sterile saline solution (0.9% NaCl) until homogeneous. Suspensions were centrifuged at 7,100 xg for 30 min at room temperature. Supernatants were carefully transferred to new tubes for alginate quantification, while the pellets were resuspended in an equal volume of LB for colony-forming unit (CFU) determination. For alginate quantification, 70 µL of each supernatant was mixed with 600 µL of freshly prepared, ice-cold 100 mM borate (SIGMA #B0394) solution in H_2_SO_4_. Samples were vortexed for 4 s and kept on ice. Subsequently, 20 µL of 0.1% (w/v) carbazole (SIGMA #442506) dissolved in absolute ethanol was added to each sample. Samples were incubated at 55°C for 30 min, and absorbance was measured at 530 nm. A calibration curve was generated using sodium alginate solution (Sigma #W201502) ranging from 50 to 500 µg/mL. All samples were analyzed in triplicate.

#### Bacterial viability assay

Overnight cultures were initiated from single colonies. The following day, cultures were diluted to an OD_600_ of 0.1 in LB supplemented with the previously specified antibiotic (5 µg/mL Aztreonam,125 µg/mL A22, and 450 µg/mL Mecillinam. Cultures were incubated at 37°C with constant shaking for 4 hours. Samples were collected at two time points: immediately after the addition of the antibiotics (t_0_) and 4 hours of incubation. At each timepoint, cultures were serially diluted (10⁻¹ to 10⁻⁷) in LB. From each dilution, 10 µL were spotted in triplicate onto LB agar plates. Plates were incubated at 30°C overnight. Experiments were performed with at least three independent biological replicates.

#### PG synthesis analysis by HADA labeling

PG biosynthesis was monitored by incorporation of the fluorescent D-amino acid HADA (7-hydroxycoumarin-3-carboxylic acid–D-alanine) (96). Overnight cultures were diluted to an OD_600_ of 0.05 and grown at 37°C to OD_600_= 0.2. Cultures were subsequently re-diluted to OD₆₀₀ = 0.05 and grown at 37°C with shaking to OD_600_= 0.4 (early exponential phase) to enrich actively dividing cells. HADA was added to a final concentration of 750 µM, and cultures were incubated for 30 min at 37°C with agitation to label nascent PG. Cells were washed twice with 1 mL PBS to remove unincorporated dye and fixed with 4% (w/v) paraformaldehyde in PBS for 1 h 45 min at 4°C. After fixation, cells were washed twice with PBS and mounted on 1.5% agarose pads for imaging. Fluorescence imaging was performed using an inverted Olympus IX83 confocal microscope equipped with a Hamamatsu ORCA-Flash4.0 sCMOS camera and a 100× objective. HADA fluorescence was excited at 385 nm. Phase-contrast images were collected to assess cell morphology and confirm bacterial presence.

Cells at different stages of division were quantified by manual counting using the ImageJ program and Cell Counter plugin (88).

Newly synthesized PG distribution was assessed by overlaying phase-contrast and HADA fluorescence channels. Cell length and width within ROIs were quantified using MicrobeJ (version 5.13p). Segmentation parameters were optimized separately for rod-shaped wild-type cells and spherical mutants.

HADA fluorescence was background-subtracted prior to analysis. Automated segmentation was manually curated, and incorrectly segmented cells were excluded.

Demographs and heat maps of HADA signal distribution along the longitudinal cell axis were generated using MicrobeJ (28). For heat map analysis, cells were categorized into four classes based on cell length.

#### PG purification

PG sacculi were purified as described in Corona and Vollmer (48). Briefly, four liters of exponentially growing bacterial culture were harvested by centrifugation (6,000 x g, 15 min). Cell pellets were resuspended in an ice-cold 50 mM MES pH 6.0 buffer. The suspension was added dropwise to a boiling 8% SDS solution under continuous stirring and incubated at 95-100°C for 1 hour to lyse the cells and solubilize proteins. Samples were kept at room temperature overnight to cool down and were then centrifuged for 20 min at 46,000 x g at room temperature. The resulting pellets containing the crude sacculi were washed repeatedly with 50 mM MES (pH 6.0). Washing was repeated at least eight times until SDS was no longer detected through the Hayashi assay (97). Briefly, 250 µL of the sample was mixed with 75 µL Hayashi reagent (0.005% methylene blue in 0.7 mM phosphate buffer, pH 7.2), followed by the addition of 750 µL of chloroform. Samples were vortexed for 20-30 s, then kept steady to allow phase separation. The blue coloration in the lower chloroform phase indicated the presence of SDS. The sacculi were considered SDS-free when the chloroform phase remained colorless, and the aqueous phase retained the dye.

The samples were resuspended in 7.5 mL of 50 mM MES buffer, pH 6.0, and treated with 200 µg of α-amylase (to degrade polysaccharide contaminants) and 300 µg of Pronase E (to remove covalently bound lipoproteins). Samples were incubated overnight at 37°C on an end-over-end rotator. Following enzymatic digestion, the sacculi were washed with the MES buffer supplemented with an EDTA-free protease inhibitor tablet. The final pellet was then resuspended in MilliQ pure water and stored at 4°C for short-term use, and at 20°C for long-term storage.

### Atomic Force Microscopy (AFM)

#### Sample preparation

The sacculi, purified as described above, were immobilized on freshly cleaved mica using Cell-Tak-mediated surface functionalization. To activate the surface, 57 µL of 1 M NaHCO₃ was deposited onto the mica. Then, 0.5 µL of Cell-Tak adhesive (2 mg/mL, Corning#354240) was added and gently spread across the surface using a pipette tip until evenly distributed. Then, 1 µL of 1 M NaOH was added and mixed carefully, avoiding any scratching of the mica surface. The activation process was allowed to proceed for 25 minutes at room temperature. The surface was then thoroughly rinsed with ultrapure water and dried with a stream of nitrogen.

For sample immobilization, 20 µL of sacculi sample (∼1 mg/mL) were deposited onto the activated mica and incubated for 20 minutes to allow binding. In some preparations, samples were additionally incubated in a vacuum desiccator to promote surface attachment. Following incubation, excess liquid was gently removed, and the surface was kept hydrated during AFM imaging by adding ultrapure water to avoid sacculi adherence to AFM probe tips and interfere with image acquisition.

#### AFM imaging

AFM was performed in liquid mode using a Bruker Multimode 8 system equipped with a Nanoscope V controller and Nanoscope 9.2 software. Images were acquired using silicon nitride cantilevers (ScanAsyst-FLUID, Bruker AFM Probes, Camarillo, CA, USA) with nominal specifications of k = 0.7 N/m, Fq = 150 kHz, and a tip radius of 20 nm. First, a quick survey of the sample was performed in tapping mode to locate the particles of interest. Then, further imaging was performed using the PeakForce QNM mode to scan the sample locally. Probe and QNM calibration were performed according to the manufacturer’s setup menu in NanoScope 9.2. Square images were acquired at around 1Hz scan rate at variable scan sizes ranging from 1 µm to 10 µm, with a sampling of 512 to 1024 pixels. Force was manually controlled within a range of 600–800 pN.

#### Image Processing

AFM images were corrected using Gwyddion software (98,99). The raw images were corrected using mean-plane subtraction and row alignment with the median method. Flattening was finalized by isolating particles above a threshold, and the background was flattened using a third-order polynomial. Large scan images were cropped to isolate the sacculus, for which threshold values were used to select the sacculus and background areas separately. Images of 2 µm and 3 µm were used to analyze the samples. Particles that overlapped one another were excluded from the analysis because it was not possible to measure their true height.

Sacculi were identified individually as grains in Gwyddion using a height threshold. The average grain heights of the sacculi and the background were measured separately to determine the relative height in Gwyddion. The average relative height value was used for the final statistics.

### Synthetic lethality

#### Transposon mutant library generation

Transposon mutant libraries were constructed as described in Janet-Maitre et al., 2023 (100). Briefly, *P. aeruginosa* CHA wild-type and *mreD^down^* cultures grown overnight at 42°C with shaking were combined with two *E. coli* strains harboring pRK2013 and pBTK24. Conjugation mixtures, consisting of 100 μL of each culture normalized to an OD_600_ of 1, were pelleted, washed in LB, spotted onto pre-warmed LB agar plates, and incubated at 37°C for 5 h. Cells were then recovered into liquid LB, and aliquots were plated on LB agar supplemented with Irgasan (25 μg/mL) and Gentamicin (75 μg/mL). After incubation at 37°C, colonies were pooled directly into LB containing 20% glycerol, aliquoted, and stored at −80°C.

#### Sample preparation

Transposon mutant libraries were thawed on ice, and 1mL of bank was inoculated into 30mL of LB, and bank growth was performed overnight at 27°C to limit growth. Overnight cultures were diluted to OD_600_=0.1 in LB and incubated at 37°C under agitation until they reached OD_600_=1. Experiments were performed in biological duplicates. Genomic DNA was extracted using GenElute bacterial genomic DNA kit (Sigma #<u>NA2110</u>).

#### Library

Libraries were constructed as described in (Janet-Maitre et al., 2021). Briefly, genomic DNA was fragmented, end-repaired, and dA-tailed (NEB), followed by ligation to annealed short and long adaptors (Table S6). Fragments of 200–400 bp were size-selected on a 2% agarose gel. Purified adaptor-DNA fragments were amplified by Phusion polymerase (NEB) in combination with specific primers (PCR1 Tn-specific and PCR1 Adaptor comp; Table S6). A second amplification round was performed with a primer bearing a P5 Illumina sequence and P7 indexed Illumina primers (NEB). After each step, products were quantified using a Qubit dsDNA HS Assay kit (Q33230). The quality of the libraries was assessed on an Agilent Bioanalyzer 2100 using high-sensitivity DNA chips.

#### Sequencing and data analysis

Libraries were sequenced on an Illumina NextSeq 500 (I2BC, Saclay Paris). Sequencing reads were trimmed and mapped to the genome of CHA (NCBI accession number GCF_003698505.1) using Bowtie2 (101). Reads were assigned to features using Ht-seq-count and differential representation of insertion mutants between two strains was performed using DESeq2 (102,103). The chord plot was generated using the Circos program on the usegalaxy.eu online platform (Galaxy Version 0.69.8+galaxy9) (104). Functional enrichment analysis was performed on the STRING database (Version 12.0; (52)) by analyzing all significant hit (|Log_2_FC| > 1; *P_adj_* < 0.05), and beneficial and detrimental hits independently using the PAO1 gene identifiers.

### Transcriptomics

#### RNA extraction

Bacteria were grown overnight at 37°C under agitation. Strains were subcultured at OD_600_=0.01 and grown to OD_600_=1, either in neutral LB (pH 6.8), LB pH8.8 (pH adjusted using NAOH 6M) or Neutral LB supplemented with carbenicillin (20µg/mL). Bacterial pellets were frozen with liquid nitrogen when they reached the appropriate OD_600_ until all the samples were ready. RNA extraction and library preparation were performed as previously described (100).

#### Sequencing and data analysis

Sequencing was performed by the I2BC sequencing facility (Saclay Paris) on an Illumina Nextseq 500 instrument. Raw data was processed and mapped onto the CHA genome (NCBI accession number GCF_003698505.1) using Bowtie2 (101). Reads were assigned to features using Htseq-count (102,103) and differential gene expression between strains was performed by DESeq2 (102,103). RNA-seq experiment was performed on biological duplicates.

#### Competition assays

For competition assays, *mreD^down^*Δ*Tc* (Tetracycline-sensitive) was transformed with pJN105 (Gm^R^). The mutant strains used: *mreD^down^* Δ*ctpA, mreD^down^* Δ*ponA*, *mreD^down^* Δ*RS11480*, *mreD^down^* Δ*minD,* and *mreD^down^* Δ*lpoP* are Tetracycline-resistant. Bacterial strains were cultured overnight in LB rich medium at 37°C with agitation. Cultures were diluted to OD_600_=0.1 to ensure active cell division. Once cultures reached an OD_600_∼1, they were used to inoculate 2 mL LB medium. Equal inocula of the wild-type strain and each mutant strain (OD_600_∼0.05) were mixed. Mixed cultures were incubated at 37°C with shaking for 24 hours. Samples were collected at two timepoints: immediately after mixing (t_0_) and after 24 hours of incubation (t_24hrs_). At each timepoint, cultures were serially diluted (10^-1^ to 10^-7^) in LB. From each dilution, 10 µL were spotted in triplicate onto LB agar, LB agar supplemented with 75 µg/mL of Tetracycline, and LB agar supplemented with 75 µg/mL of Gentamicin agar square plates. Plates were incubated at 37°C overnight. Experiments were performed with five independent biological replicates. The next morning, wild-type and mutant strains were enumerated, and the competitive index was calculated using the following formula: (t_24hrs_ mutant CFU/t_24hrs_ wild-type CFU)/(t_0_ mutant CFU/t_0_ wild-type CFU).

#### Statistical analysis

Details on the number of replicates, sample sizes, and statistical tests are given in the text or figure legends. All statistical analyses were performed using GraphPad Prism software.

## Data availability

The data generated for Tn-seq and RNA-seq have been deposited on Gene Expression Omnibus (GEO) under the accession number GSE325015 and GSE325145, respectively. The raw whole-genome sequencing data was deposited on the Sequence Read Archive (SRA) under the bioproject name PRJNA1445359.

## Acknowledgement

The work was supported by grants from Agence Nationale de la Recherche (C7H-ANR23A43), the Laboratory of Excellence GRAL, financed within the University Grenoble Alpes graduate school (Écoles Universitaires de Recherche) CBH-EUR-GS 785 (ANR-17-EURE-0003). The sequencing part has benefited from the facilities and expertise of the high-throughput sequencing core facility of I2BC (Centre de Recherche de Gif – http://www.i2bc.paris-saclay.fr/). This work used M4D facilities at the Grenoble Instruct-ERIC Center (ISBG; UMS 3518 CNRS CEA-UGA-EMBL) with support from the French Infrastructure for Integrated Structural Biology (FRISBI; ANR-10-INSB-05-02) and GRAL, a project of the University Grenoble Alpes graduate school (Ecoles Universitaires de Recherche) CBH-EUR-GS (ANR-17-EURE-0003) within the Grenoble Partnership for Structural Biology. IBS acknowledges integration into the Interdisciplinary Research Institute of Grenoble (IRIG, CEA). We also acknowledge the financial support of the Cross-Disciplinary Program on Instrumentation and Detection (PTC-ID LCTEM) of CEA, the French Alternative Energies and Atomic Energy Commission. M. JM received a Ph.D. fellowship from the French Ministry of Education and Research. The authors thank Mylène Robert-Genthon and Aimé Berwa for their help in the construction of WGS libraries. We also thank Stephanie Bouillot for her help in the image analysis and immunofluorescence experiments, and Adrien Ducret for his help in MicrobeJ analysis.

