## Supplementary Figures 1-7 and Supplementary tables 3-6 for "Elongasome Dysfunction Triggers Dependence on MepM-Mediated Peptidoglycan Recycling"

[1] Univ. Grenoble Alpes, CEA, CNRS, IBS, Team Bacterial Pathogenesis and Cellular Responses, F-38000 Grenoble, France.

[2] Univ. Grenoble Alpes, CEA, CNRS, IBS, F-38000 Grenoble, France.

* Current address: Institute for Molecular Infection Biology (IMIB), Faculty of Medicine, University of Würzburg, Würzburg, Germany

[#] Authors contributed equally to the work

### Supplementary information

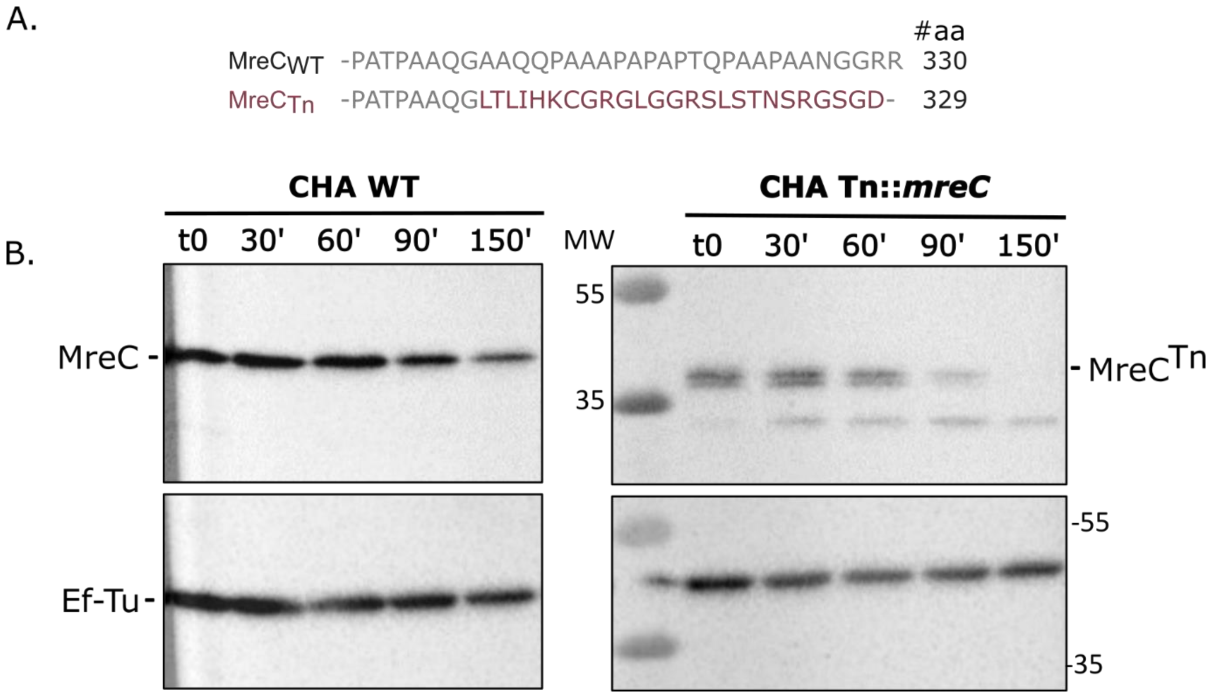

**Supplementary Figure 1. Transposon insertion at the end of *mreC* gene affects MreC protein abundance and stability. A.** The C-terminal region of MreC protein sequence in CHA wild-type (WT) and Tn::*mreC* at the transposon insertion site. **B.** Culture of CHA WT and transposon mutant were grown in LB. Samples were collected every 30 min. MreC prodution was followed by immunoblot on total bacteria using anti-MreC antibodies. Anti-Ef-Tu was used as loading control. MW: Molecular Weight (kDa).

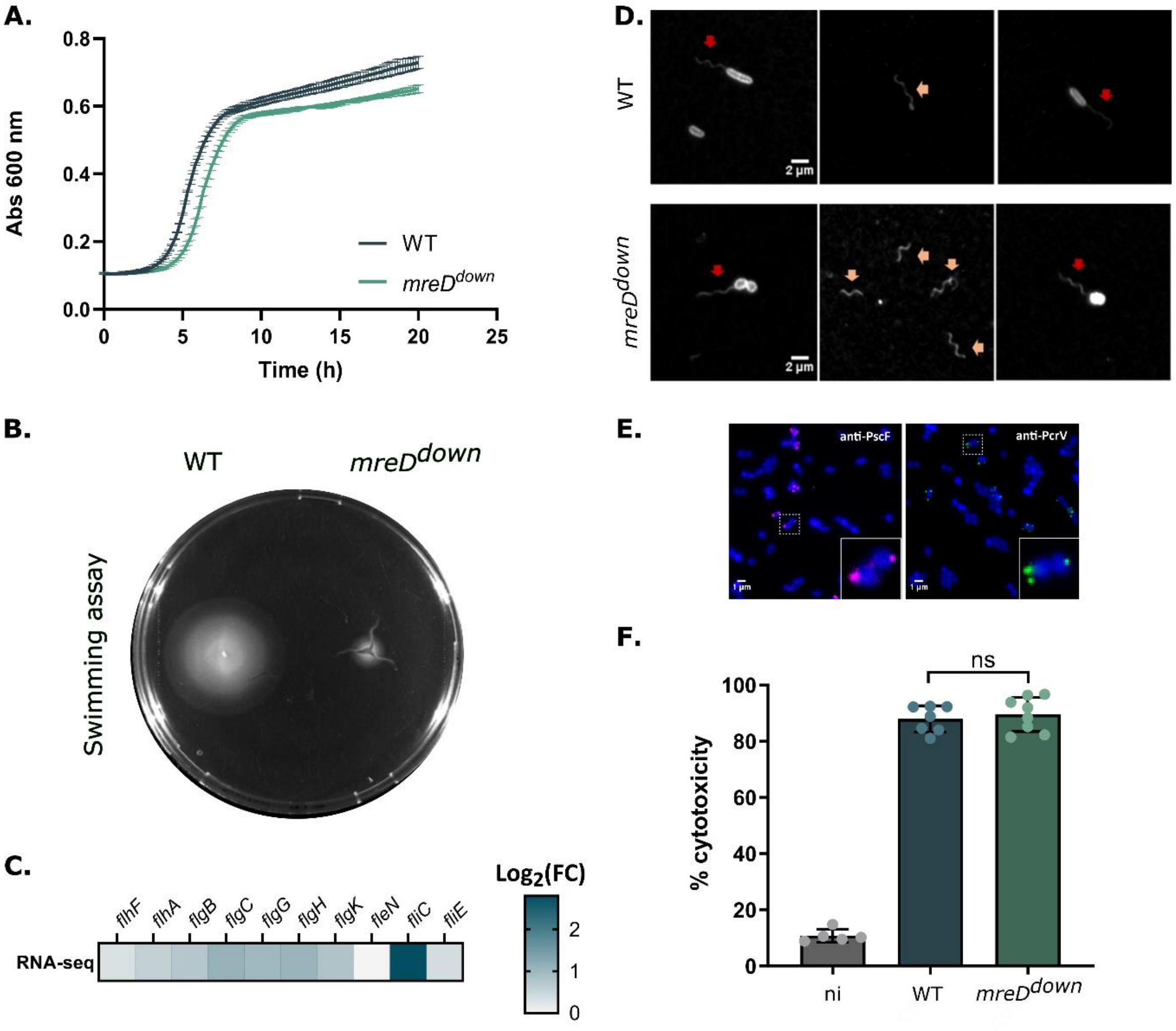

**Supplementary Figure 2: Downregulation of *mreD* alters bacterial motility but not cytotoxicity. A.** Growth of the wild-type (WT) and *mreD*$^{down}$ strains in LB medium was monitored for 20 hours. **B.** Swimming motility of the WT and *mreD*$^{down}$ strains. **C.** Heatmap representation of transcriptomic data in flagellar genes (*mreD*$^{down}$ vs WT). **D.** FliC was visualized by immunofluorescence on fixed bacteria using anti-FliC antibody. Red arrows: flagellated cells. Orange arrows: detached flagella. **E.** T3SS components [PscF (left) and PcrV (right)] were visualized by immunofluorescence on fixed *mreD*$^{down}$ bacteria using anti-PscF and anti-PcrV antibodies. **F.** WT and *mreD*$^{down}$ cytotoxicity toward J774.A1 cells by LDH measurement. Each point represents an individual measurement, bars height represents the mean and horizontal lines indicate standard deviation. Statistical significance was assessed using an unpaired t-test. ni: not infected**.**

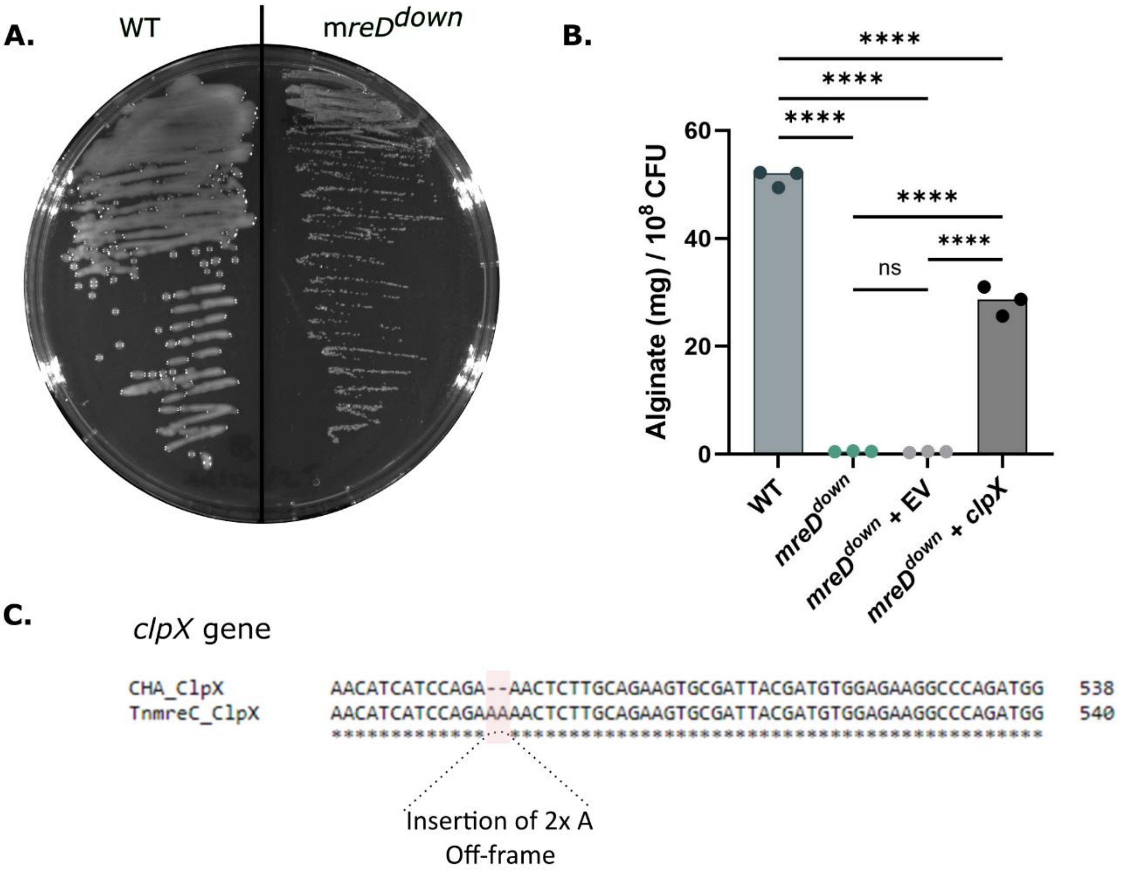

**Supplementary Figure 3. Loss of mucoidy in the *mreD*$^{down}$ strain due to an inactivating mutation in *clpX*. A.** Wild-type and *mreD*$^{down}$ strains plated on LB agar showing reduced mucoidy in the mutant strain. **B.** Alginate production was quantified using carbazole assay. Normality of the data was tested by a Shapiro-Wilk test and a one-way ANOVA was performed with Tukey's multiple comparisons test. ****: $P_{val}$<0.0001. The histogram shows the median. **C.** Sequencing revealed an insertion in the *clpX* gene that causes a frameshift, resulting in inactivation of ClpX.

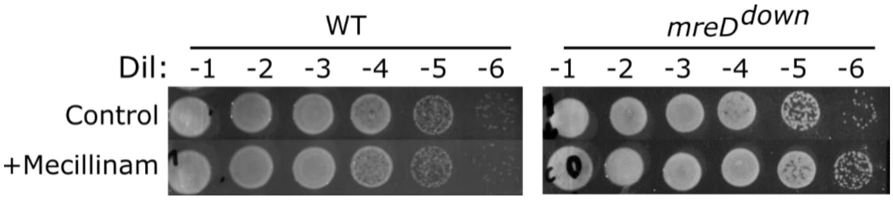

**Supplementary Figure 4: The wild-type and *mreD*$^{down}$ strains resist to high concentrations of Mecillinam.** Wild-type (WT) and *mreD*$^{down}$ were treated with 450 µg/mL Mecillinam. After 4 hours of incubation, cultures were serially diluted and spotted onto LB agar. Dil: dilution.

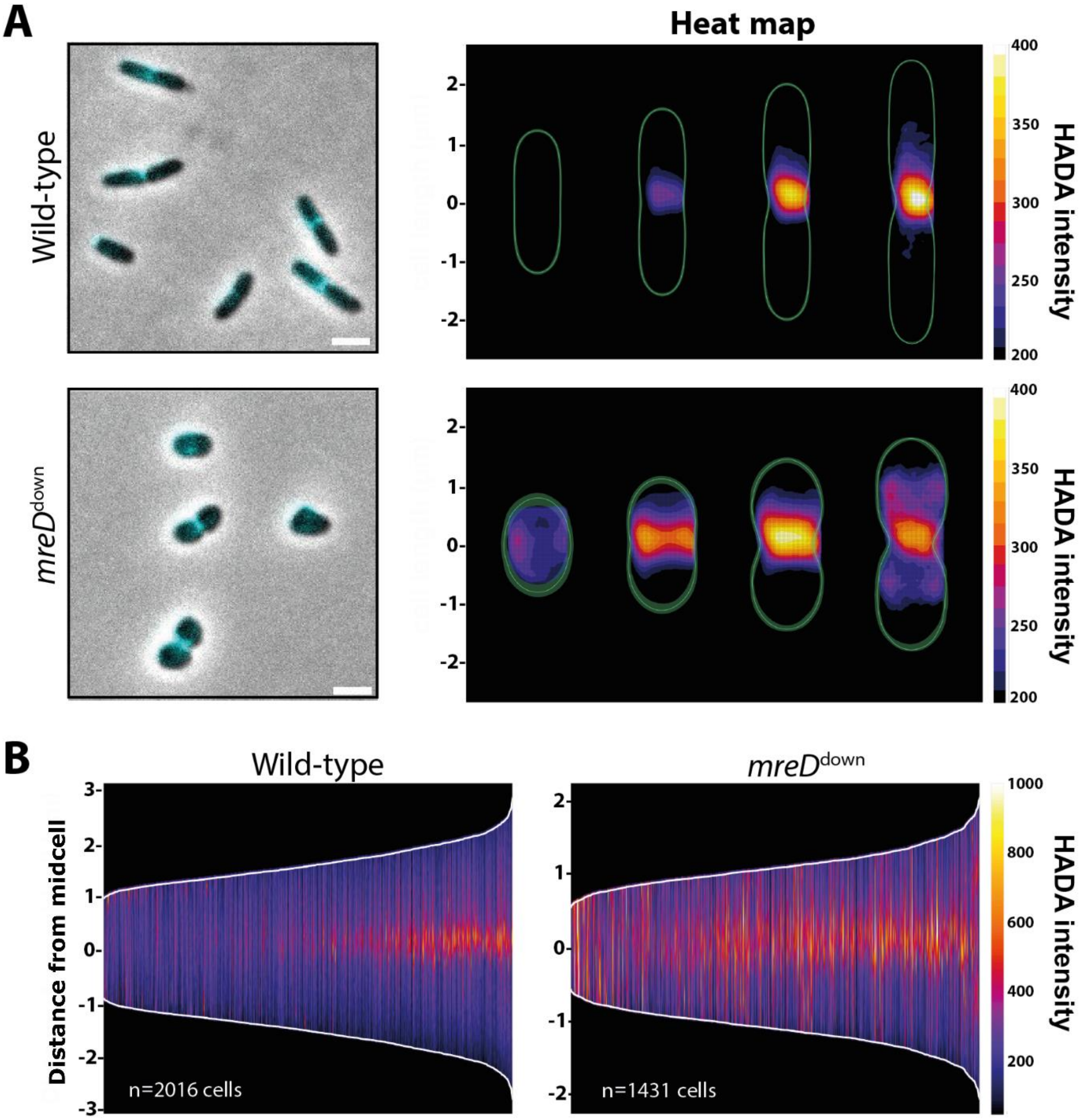

**Supplementary Figure 5: HADA incorporation reveals increased septal peptidoglycan synthesis in the mutant. A.** Representative merged images of HADA labeling and phase-contrast channels (left), with corresponding heat maps showing HADA fluorescence intensity across four cell-length classes (right). Cells within each class are aligned along their long axis and normalized to cell length to enable comparison across populations. All images were scaled identically to facilitate direct comparison. Scale bar: 2 µm **B.** Demographs showing the distribution of HADA fluorescence across single cells sorted by cell length, highlighting distinct stages of the cell cycle. Fluorescence intensity is mapped along a normalized cell axis centered at midcell, allowing comparison of spatial patterns of peptidoglycan synthesis. Septal peptidoglycan synthesis is more pronounced at all growth stages in the *mreD*$^{down}$ mutant compared to the wild type.

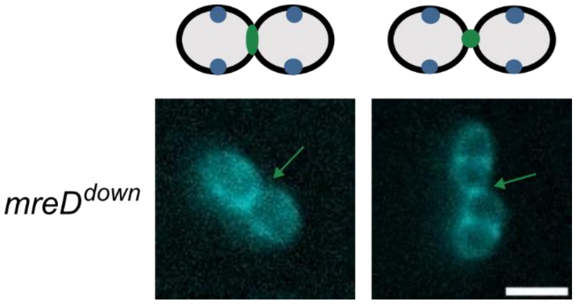

**Supplementary Figure 6: *mreD*$^{down}$ initiates division before completion of the previous cycle.** HADA labeling reveals sites of new PG synthesis. Septa represented in blue indicate newly initiated division sites, whereas septa in green correspond to septa from the previous division cycle (green arrow). Scale bar: 2 µm

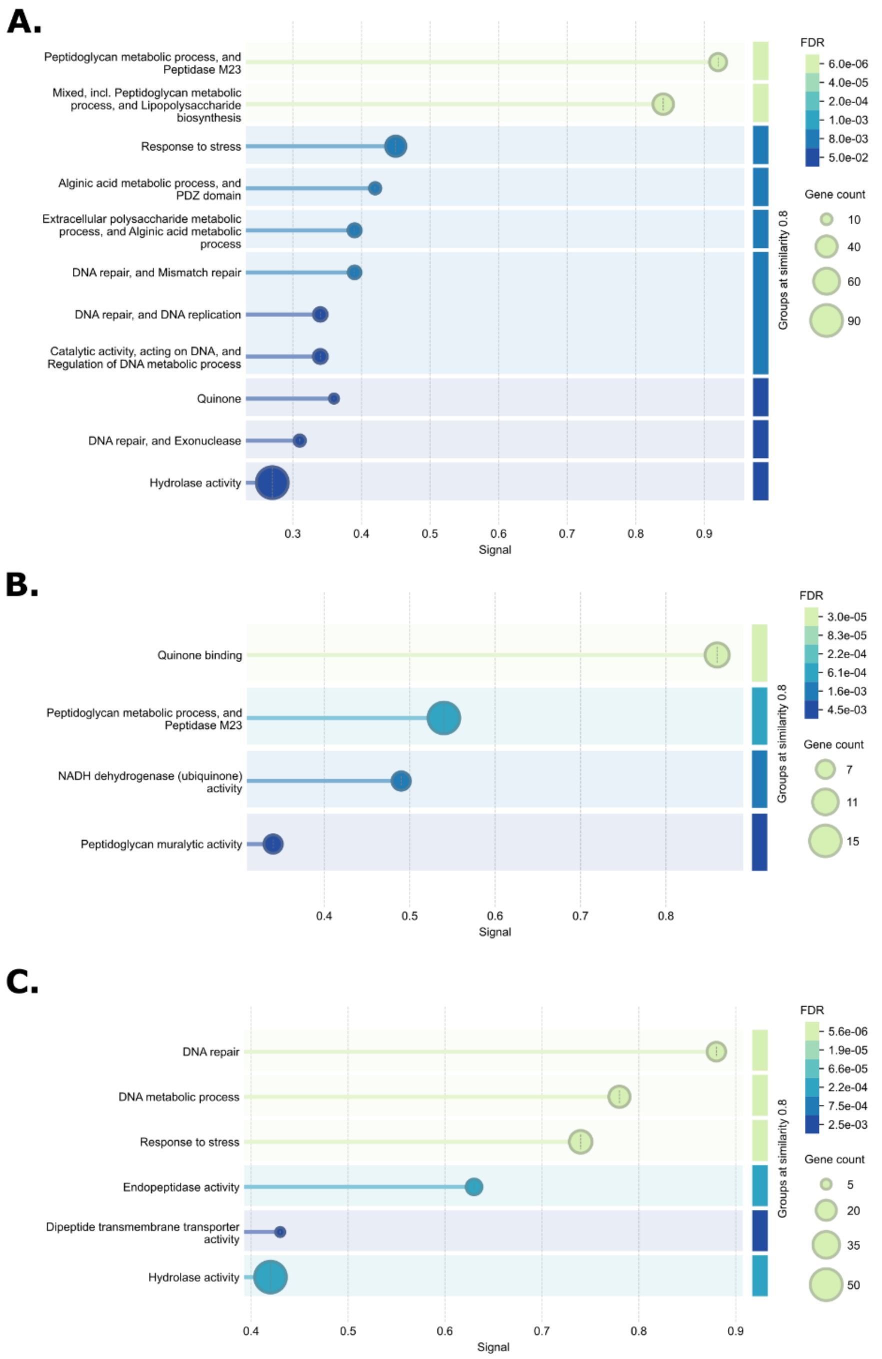

**Supplementary Figure 7. Functional enrichment analysis of the synthetic lethal data. A.** Functional enrichment analysis of all significant hits ($|Log_2(FC)|>1$, $P_{adj}<0.05$). **B.** Functional enrichment analysis of all beneficial hits ($Log_2(FC)>1$, $P_{adj}<0.05$). **C.** Functional enrichment analysis of all detrimental hits ($Log_2(FC)<1$, $P_{adj}<0.05$). Plots were generated by STRING (1).

**Supplementary table 1. Differential gene expression in various growth conditions**

**Supplementary table 2. Synthetic lethality screen dataset**

**Supplementary table 3. Genes whose disruption results in synthetic lethality* in the *mreD*$^{down}$ background**

* Synthetic lethality was determined when less than 50 reads were detected on average across *mreD*$^{down}$ replicates. # Gene identifiers (ID) (NCBI accession number GCF_003698505.1 and their PAO1 homolog). 'IPC1262_' has been removed from all identifiers for readability.

| CHA_ID# | PAO1_ID | Name | Product Name | $Log_2$(FC) | $P_{adj}$ |
|---|---|---|---|---|---|
| *RS28395* | *PA0762* | *algU* | sigma factor AlgU | -9.48 | 5.51E-93 |
| *RS02905* | *PA3999* | *dacC* | D-ala-D-ala-carboxypeptidase | -4.36 | 2.35E-36 |
| *RS13795* | *PA0596* | *amgK* | hypothetical protein | -4.29 | 1.48E-21 |
| *RS13545* | *PA0663* | | hypothetical protein | -4.32 | 5.37E-21 |
| *RS13525* | *PA0667* | *mepM* | MepM | -5.50 | 1.56E-19 |
| *RS21305* | *PA2329* | | probable ATP-binding component of ABC transporter | -6.60 | 9.09E-18 |
| *RS15480* | *PA0298* | *spuB* | Glutamylpolyamine synthetase | -3.50 | 8.02E-17 |
| *RS22800* | *PA4699* | *lpoP* | hypothetical protein | -6.01 | 1.30E-15 |
| *RS14415* | *PA0501* | *bioF* | 8-amino-7-oxononanoate synthase | -5.81 | 3.00E-14 |
| *RS23145* | *PA3244* | *minD* | cell division inhibitor MinD | -5.71 | 2.14E-13 |
| *RS13790* | *PA0597* | *murU* | probable nucleotidyl transferase | -5.31 | 3.05E-12 |
| *RS14905* | *PA0406* | *tonB3* | TonB3 | -3.37 | 6.61E-12 |
| *RS27165* | *PA4422* | *yraL* | conserved hypothetical protein | -4.79 | 7.42E-11 |
| *RS27385* | *PA4381* | *colR* | two-component response regulator ColR | -5.50 | 1.35E-10 |
| *RS13120* | *PA3172* | *mupP* | probable hydrolase | -5.75 | 4.07E-10 |
| *RS29195* | *PA4553* | *pilX* | type 4 fimbrial biogenesis protein PilX | -4.36 | 2.44E-09 |
| *RS25220* | *PA5562* | *parB* | chromosome partitioning protein Spo0J | -4.06 | 4.23E-09 |
| *RS27145* | *PA4426* | *yraP* | conserved hypothetical protein | -3.28 | 5.99E-08 |
| *RS02040* | *PA3832* | *holC* | DNA polymerase III, chi subunit | -4.39 | 5.09E-07 |
| *RS11630* | *PA2884* | | hypothetical protein | -3.58 | 6.70E-07 |
| *RS07620* | *PA0964* | *pmpR* | pqsR-mediated PQS regulator, PmpR | -3.33 | 2.71E-06 |
| *RS27390* | *PA4380* | *colS* | two-component sensor ColS | -3.85 | 4.63E-06 |
| $P_{RS06480}$ | *PA1064* | | hypothetical protein | -4.17 | 7.22E-05 |
| *RS14405* | *PA0503* | *bioC* | probable biotin synthesis protein BioC | -3.69 | 1.96E-04 |
| *RS13900* | *PA0576* | *rpoD* | sigma factor RpoD | -3.52 | 2.14E-04 |
| $P_{RS31020}$ | *PA4268* | $P_{rpsL}$ | 30S ribosomal protein S12 | -3.91 | 6.11E-04 |
| $P_{RS18815}$ | *PA1529* | $P_{lig}$ | DNA ligase | -3.14 | 1.07E-03 |
| *RS23260* | *PA3263* | *rdgC* | recombination-associated protein RdgC | -3.15 | 1.08E-03 |
| *RS13240* | *PA4222* | | probable ATP-binding component of ABC transporter | -3.06 | 2.34E-03 |
| *RS01560* | *PA3752* | | hypothetical protein | -3.16 | 2.77E-03 |
| $P_{RS09950}$ | *PA5038* | $P_{aroB}$ | 3-dehydroquinate synthase | -3.03 | 1.19E-02 |
| $P_{RS00960}$ | *PA3634* | $P_{ybgQ}$ | conserved hypothetical protein | -3.18 | 1.19E-02 |

### Supplementary table 4. Tn-seq data for *P. aeruginosa* endopeptidases

| CHA_ID | PAO1_ID | Name | $Log_2$(FC) | $P_{adj}$ |
|---|---|---|---|---|
| IPC1262_RS13525 | PA0667 | mepM | -5.50 | 1.56E-19 |
| IPC1262_RS27260 | PA4404 | | -3.21 | 1.75E-15 |
| IPC1262_RS01785 | PA3787 | | -1.86 | 1.59E-02 |
| IPC1262_RS00905 | PA3623 | | -0.16 | 6.57E-01 |
| IPC1262_RS10560 | PA4924 | | 0.39 | 2.43E-01 |
| IPC1262_RS25165 | PA5551 | | -1.01 | 7.41E-04 |

### Supplementary table 5. Bacterial strains and plasmids

| **Bacteria** | **Features / Source** | **Reference/origin** |
|---|---|---|
| *Pseudomonas aeruginosa* | | |
| CHA | T3SS+ , ExoS+, Cystic fibrosis isolate | Toussaint et al, 1993 (2), Bezuidt 2013 (3) |
| *mreD*$^{down}$ (CHA::Tn*mreC)* | | Dacheux 2002 (4) |
| *mreD*$^{down}$ΔTcR | | This work |
| *mreD*$^{down}$/pJN105 | | This work |
| *mreD*$^{down}$/pJN105-*mreC* | | This work |
| *mreD*$^{down}$/pJN105-RBS$_{tssK}$*mreC* | | This work |
| *mreD*$^{down}$/pJN105-RBS$_{tssK}$*mreD* | | This work |
| *mreD*$^{down}$/pJN105-RBS$_{tssK}$*mreCD* | | This work |
| CHA/pJN105 | | This work |
| CHA/pJN105-RBS$_{tssK}$*mreC* | | This work |
| CHA/pJN105-RBS$_{tssK}$*mreD* | | This work |
| CHA/pJN105-RBS$_{tssK}$*mreCD* | | This work |
| *mreD*$^{down}$ΔTcR/pSW196 | | This work |
| *mreD*$^{down}$ΔTcR/pSW196-*clpX* | | This work |
| *mreD*$^{down}$ΔTcR/pSW196-*mepM-mCherry* | | This work |
| *mreD*$^{down}$ΔTcR/pSW196-*ctpA-mCherry* | | This work |
| *Escherichia coli* | | |
| DH5α | Laboratory strain | Lab collection |
| TOP10 | Cloning strain | Invitrogen |
| **Plasmid** | **Features / Source** | **Reference/origin** |
| pBTK24 | Plasmid with Himar-1 mariner transposon and C9 transposase (Amp$^R$, Gm$^R$) | Kulasekara *et al*, 2004 (5) |
| pRK600 | Helper plasmid with conjugative properties (Cm$^R$) | Kessler *et al*, 1992 (6) |
| pJN105 | Plasmid for protein expression in *Pseudomonas* inducible by arabinose (Gm$^R$). Used for complementation of Tn::*mreC* | This work |

| | | |
|---|---|---|
| pJN105-*mreC* | Plasmid expressing *mreC* from CHA strain | This work |
| pJN105-RBS$_{tssK}$-*mreC* | Plasmid expressing *mreC* from CHA strain with the RBS of TssK | This work |
| pJN105-RBS$_{tssK}$-*mreD* | Plasmid expressing *mreD* from CHA strain with the RBS of TssK | This work |
| pJN105-RBS$_{tssK}$-*mreCD* | Plasmid expressing *mreC* and *mreD* from CHA strain with the RBS of TssK | This work |
| pSW196 | Integrative plasmid in *attB* site, used for protein expression in *Pseudomonas* inducible by arabinose ($Tc^R$) | This work |
| pSW196-*clpX* | Plasmid expressing *clpX* from CHA | This work |
| pSW196-*mepM-mCherry* | Plasmid expressing *mepM* from CHA with an mCherry fluorescent tag at its C-terminus | This work |
| pSW196-*ctpA-mCherry* | Plasmid expressing *ctpA* from CHA with an mCherry fluorescent tag at its C-terminus | This work |
| pEXG2 | Suicide vector for allelic exchange in *Pa* ($Gm^R$)- *sacB* | This work |
| pEXG2-KO-*clpX* | pEXG2 with flanking regions to delete *clpX* | This work |
| pEXG2-KO-*ponA* | Suicide vector to delete *ponA* | This work |
| pEXG2-KO-*mepM* | Suicide vector to delete *mepM* | This work |
| pEXG2-KO-*mrcB* | Suicide vector to delete *mrcB* | This work |
| pEXG2-KO-*ctpA* | Suicide vector to delete *ctpA* | This work |
| pEXG2-KO-*minD* | Suicide vector to delete *minD* | This work |
| pEXG2-KO-*lpoP* | Suicide vector to delete *lpoP* | This work |
| pEXG2-KO-*PA2854* | Suicide vector to delete *PA2854* | This work |
| pEXG2-KO-ΔTcR | pEXG2 with flanking regions to delete Tetracycline cassette | This work |

**Supplementary table 6. Oligonucleotides used for PCRs**

| Primers | Sequence (5'-3') | |
|---|---|---|
| Eco-RBS-MreC | AAAGAATTCAAAGGCCCGTCGCTGGG | Overexpression of *mreC* gene in pJN105-*mreC* |
| MreC-SacI | TTTGAGCTCTCAGCGGCGCCCCCCGT | Overexpression of *mreC* gene in pJN105-*mreC* |
| RBS_TssK_NdeI_rw | CCCGTTTTTTTGGGCTAGCGAATTCACCTTCGGAGTCCCATATGCTGGCCGTGCTGTCGGTCAGCC | Substitution of $RBS_{mreC}$ with the $RBS_{tssK}$ |
| RBS_TssK_NdeI_fw | GGCTGACCGACAGCACGGCCAGCATATGGGACTCCGAAGGTGAATTCGCTAGCCCAAAAAAACGGG | Substitution of $RBS_{mreC}$ with the $RBS_{tssK}$ |
| NdeI-MreD | AAACATATGGCTCGCGCGACCC | Overexpression of *mreD* gene in pJN105-*mreD* |
| MreD-SacI | TTTGAGCTCCCTACCTCACGTTGAGACGCAGG | Overexpression of *mreD* and mreCD |
| NdeI-MreC | AAACATATGCTGGCCGTGCTGTCGG | Overexpression of mreCD pJN105-*mreCD* |
| sF1-pSW196-*clpX*-comp | GCTAGCGAATTCCTGCAGCCCCCTTCATCTTGTGTTTGAAGCT | Overexpression of *clpX* in pSW196-*clpX* |
| sR1-pSW196-*clpX*-comp | TCTAGAACTAGTGGATCCCCCCACGAAACAGATAGGAAGAC | Overexpression of *clpX* in pSW196-*clpX* |
| F1-pSW196-*mepM*-mCherry-comp | CCACTAGTACCTTCGGAGTCCCATAAAGCTTGTGTTCCCCTCGAGCGAAGTC | Overexpression of *mepM-mCherry* in pSW196-*mepM-mCherry* |
| R1-pSW196-*mepM*-mCherry-comp | CTCACCATGCCGCCGGCGGCGGC AAAGCTTGTTGGCCCTGAACAAGCAGCGC | Overexpression of *mepM-mCherry* in pSW196-*mepM-mCherry* |
| F1-pSW196-*ctpA*-mCherry-comp | CCACTAGTACCTTCGGAGTCCCATAAAGCTTATGCTGCATTGCTTCCGTCCC | Overexpression of *ctpA-mCherry* in pSW196-*ctpA-mCherry* |
| R1-pSW196-*ctpA*-mCherry-comp | CTCACCATGCCGCCGGCGGCGGCAAGCTTGTTGCCGCGGGTGACGCTC | Overexpression of *ctpA-mCherry* in pSW196-*ctpA-mCherry* |
| F1-KO-*ponA* | GGTCGACTCTAGAGGATCCCCGACCACGGCCGACTTCAGG | *ponA* deletion |
| R1-KO-*ponA* | CGGTCACCGGGTCGATGCtcaGAGGGGGATCTGCAGCTGG | *ponA* deletion |
| F2-KO-*ponA* | GCATCGACCCGGTGACCG | *ponA* deletion |
| R2-KO-*ponA* | ACCGAATTCGAGCTCGAGCCCCAACAACGTGCTCGGTTTCCC | *ponA* deletion |
| F1-KO-*mrcB* | GGTCGACTCTAGAGGATCCCCCTCCAGGTAGAGATCGTCG | *mrcB* deletion |
| R1-KO-*mrcB* | GTTCTGGCTCCCGCAGCCGAGGCCGAGCTTGAGAGCC | *mrcB* deletion |
| F2-KO-*mrcB* | GGCTGCGGGAGCCAGAAC | *mrcB* deletion |
| R2-KO-*mrcB* | ACCGAATTCGAGCTCGAGCCCTCCAACTGCTCGTCGGCCG | *mrcB* deletion |
| F1-KO-*ctpA* | GGTCGACTCTAGAGGATCCCCTGCCGTGGCCGGTGAATG | *ctpA* deletion |

| | | |
|---|---|---|
| R1-KO-*ctpA* | CGCTCAGGCCTTTCAGCAGCAGCGCCAGGGTGGTGGG | *ctpA* deletion |
| F2-KO-*ctpA* | CTGCTGAAAGGCCTGAGCG | *ctpA* deletion |
| R2-KO-*ctpA* | ACCGAATTCGAGCTCGAGCCCAGTTCCTCGGCGAGCCAGG | *ctpA* deletion |
| F1-KO-*minD* | GGTCGACTCTAGAGGATCCCCGAAGAGAAGCCCGCCGAGC | *minD* deletion |
| R1-KO-*minD* | CACATCGAGGAATCGATGCGGTGGTTTTACCGACGCCAC | *minD* deletion |
| F2-KO-*minD* | CCGCATCGATTCCTCGATGTG | *minD* deletion |
| R2-KO-*minD* | ACCGAATTCGAGCTCGAGCCCACCAGCAGGGCGTCATC | *minD* deletion |
| F1-KO-*lpoP* | GGTCGACTCTAGAGGATCCCCGGCACCAGCAACGACTCG | *lpoP* deletion |
| R1-KO-*lpoP* | GCGGAGTCTCCCTGCTTCTCGCCCGCGAGCAGGGCACTG | *lpoP* deletion |
| F2-KO-*lpoP* | GAGAAGCAGGGAGACTCCGC | *lpoP* deletion |
| R2-KO-*lpoP* | ACCGAATTCGAGCTCGAGCCCTCTCCGCCTTGCTTGAAGC | *lpoP* deletion |
| F1-KO-ΔTc | GAAGCATAAATGTAAAGCAAGCTTTGCGGATGTTGCGATTACTTCG | Tetracycline cassette removal in Tn |
| R1-KO-ΔTc | GCACGCCATAGTGACTGGCG | Tetracycline cassette removal in Tn |
| F2-KO-ΔTc | CGCCAGTCACTATGGCGTGCCCATTATCGCCGGCATGGCG | Tetracycline cassette removal in Tn |
| R2-KO-ΔTc | GAAATTAATTAAGGTACCGAATTCGTTCTGCCAAGGGTTGGTTTGC | Tetracycline cassette removal in Tn |
| qPCR-*rpoD*_F | CTGCCGGAGGATATTTCAGA | Used for RT-qPCR |
| qPCR-*rpoD*_R | ATACGTTGATCCCCATGTCG | Used for RT-qPCR |
| qPCR-*mreB*-F1 | TGTCTACGCCGAATCCGTCC | Used for RT-qPCR |
| qPCR-*mreB*-R1 | CCGATCAGGCTGCCGTAGTT | Used for RT-qPCR |
| qPCR-*mreC*-F1 | ACGCCCGGTTCGACTATCTG | Used for RT-qPCR |
| qPCR-*mreC*-R1 | ATTTCCGCCAGGCCGTAGAA | Used for RT-qPCR |
| qPCR-*mreD*-F1 | TCACCTTCCTCGTGCTGTCC | Used for RT-qPCR |
| qPCR-*mreD*-R1 | GTAGACCACCACCAGCACCA | Used for RT-qPCR |
| Primers for Tn-seq | | |
| Short adpator | TACCACGACCA-NH2 | |
| Long adaptor | GTGACTGGAGTTCAGACGTGTGCTCTTCCGATCTGGTCGTGGTAT | |
| PCR1 Tn-specific | CACAGGAAACAGGACTCTAGAGG | |
| PCR2 adaptor complementary | GTGACTGGAGTTCAGACGTGTG | |
| P5+ Illumina | AATGATACGGCGACCACCGAGATCTACACTCTTTCCCTACACGACGCTCTTCCGATCTCTAGAGACCGGGGACTTATCAGC | |
| P7-index | CAAGCAGAAGACGGCATACGAGATNNNNNN | |
